# Mitotic entry is controlled by the plant-specific phosphatase BSL1 and cyclin-dependent kinase B

**DOI:** 10.1101/2025.07.24.666638

**Authors:** Frej Tulin, Yalikunjiang Aizezi, Andres V. Reyes, Yuji Fujieda, Arthur Grossman, Shou-ling Xu, Masayuki Onishi, Farhah Assaad, Zhi-Yong Wang

## Abstract

Cell cycle regulation is well understood in opisthokonts (fungi and metazoans) but not in plants and Apicomplexa, as some cell cycle regulators are not conserved^1–3^. In opisthokonts, cell cycle progression requires dephosphorylation of cyclin-dependent kinase (CDK) by the CDC25 phosphatase^4^. Plants have no CDC25, and thus their mechanisms of cell cycle regulation remain elusive^1,5,6^. Here, we show that the BSL1 phosphatase dephosphorylates CDKB1 to promote mitotic entry in *Chlamydomonas.* Mutations of BSL1 or CDKB1 block mitotic entry after DNA replication. BSL1 shows dynamic localization through the cell cycle at the basal bodies, spindle poles, and cleavage furrow. CDKB1 is hyperphosphorylated at T14 and Y15 residues in the *bsl1* mutant and in wild-type cells treated with DNA replication inhibitors. BSL1 binds to CDKB1 and dephosphorylates CDKB1 pT14/pY15 *in vitro*. Phospho-mimicking mutations of T14/Y15 inactivate CDKB1 function, whereas phospho-blocking mutations cause sensitivity to DNA replication inhibitors, which delay cytokinesis in wild-type cells more than cells expressing unphosphorylatable mutant CDKB1. These results indicate that CDKB1 T14/Y15 is phosphorylated to block mitotic entry before DNA replication is complete, and BSL1 dephosphorylates CDKB1 to promote mitosis. Our study demonstrates that BSL1, a phosphatase conserved in plants and Apicomplexa but absent in fungi and animals, is a CDKB1-activating mitosis-promoting factor that has evolved additional signaling functions in receptor kinase pathways in higher plants.

**One-Sentence Summary:** BSL1 is a mitosis-promoting phosphatase that activates CDKB1 in plants.

## Main Text

Progression through the eukaryotic cell cycle is highly regulated to ensure faithful propagation of the genetic material to daughter cells^7^. The mechanisms of cell cycle regulation have been studied extensively in fungal and animal model systems, which represent the opisthokont branch of eukaryotes, but remain poorly understood in other branches of eukaryotes including plants and medically significant parasites belonging to the Apicomplexa phylum^1,8^. In both fungi and metazoans, orderly progression through sequential cell cycle events is orchestrated by cyclin-dependent kinase (CDK), whose activity is regulated by not only the rising and falling level of cyclin but also checkpoint-controlled phosphorylation^4^. CDK is inactivated due to phosphorylation of threonine 14 (T14) and tyrosine 15 (Y15) residues to prevent cell cycle progression when DNA is damaged or incompletely replicated^9,10^. Dephosphorylation of pT14 and pY15 by the CDC25 phosphatase is required to activate CDK and recover from checkpoint arrest^4,9–12^. Dysfunctions in CDC25 cause defects in mitotic entry control, leading to genome instability and cancers^13^. This core CDK-CDC25 cell cycle regulation mechanism is conserved in fungi and animals. However, CDC25 is absent in plants and Apicomplexa^8,14^, which include plastid-containing parasites causing major human diseases such as malaria. How cell cycle progression is regulated without CDC25 in plants and Apicomplexan parasites remains a major mystery in biology.

Plants have evolved a plant-specific family of CDK, named CDKB, in addition to CDKA, the ortholog of fungal/animal Cdk1/Cdc2^15–18^. Genetic studies indicate that CDKA plays a dominant role in G1/S transition and CDKB1 is essential for mitotic entry^16,19,20^, in contrast to a single CDK acting in both transitions in yeast^21^. How checkpoint signaling regulates CDKA and CDKB1 activities, and thereby, transitions into S and M phases in plants, is not fully understood^6^. The T14 and Y15 residues are conserved in CDKA and CDKB1 but their functions remain elusive. Mutating T14/Y15 of Arabidopsis CDKA suggested that their phosphorylation inactivates CDKA but is not required for normal response to DNA replication stress^5^. The function of CDKB1 T14/Y15 phosphorylation has not been reported. Without CDC25 and functional evidence for importance of CDK phosphorylation during replication stress, recent research has focused on transcriptional and protein-protein interaction mechanisms that modulate CDK activities in plants^6^.

With multiple technical advantages over multicellular higher plants, the unicellular green alga *Chlamydomonas reinhardtii* is a powerful model system for genetic studies of cell division^22^. Screening for temperature-sensitive mutants identified conserved eukaryotic cell cycle regulators and plant-specific genes required for mitosis in Chlamydomonas^18,23^. These include a mutation in the BSL1 phosphatase gene that blocks mitosis after DNA replication, a phenotype similar to the *cyclin b* (*cycb*) and *cyclin-dependent kinase b* (*cdkb*) mutants^24^. The BSL1 family of proteins is characterized by an N-terminal Kelch repeat domain and a C-terminal phosphatase domain, and is thus also named protein phosphatase with Kelch-like domain (PPKL). BSL1 is universally conserved throughout the green plant kingdom and apicomplexan parasites but is absent from fungi and metazoan^25–27^. Arabidopsis contains four BSL family members, BSU1 (bri1 Suppressor1) and BSL1, 2, 3 (BSU1-like 1, 2, 3), which have been studied extensively as components of receptor kinase pathways that regulate brassinosteroid promotion of growth^28,29^, immune signaling^30^, and stomatal development^31–33^. However, the *Arabidopsis* triple mutants lacking BSL1, 2, 3 are embryonic lethal^25^, whereas *Chlamydomonas* contains a single BSL1 paralog that is essential for mitosis^18^, suggesting that the BSL members share an ancestral and essential role in *Chlamydomonas* and higher plants. Here, we investigate the function of BSL1 in cell cycle regulation in Chlamydomonas. Our results indicate that mitotic entry is arrested by phosphorylation of CDKB1 T14/Y15 and promoted by BSL1-mediated dephosphorylation.

### The BSL1 phosphatase is essential for mitosis in Chlamydomonas

To understand the function of BSL1 in mitosis, we characterized the cell cycle phenotypes of various loss-of-function alleles of *bsl1* mutants. The Chlamydomonas *bsl1-1* mutant carries a missense L46P mutation in the N-terminal Kelch domain of BSL1 and is inviable at 33°C, a temperature that does not affect the growth of wild-type cells. The phenotype of *bsl1-1* was rescued by a full-length genomic sequence of *Chlamydomonas* BSL1 (Cre01.g050850) fused with a C-terminal mVenus-3xFLAG tag (BSL1-VF; Fig. 1a), but not by a variant protein with a mutation in a conserved phosphatase active site residue (BSL1^D619N^-VF) ^34^ (Fig. 1b; Extended Data Fig. 1), indicating that the phosphatase activity is essential for BSL1 function. We disrupted the native BSL1 locus (*bsl1-ko*) using CRISPR-Cas9-guided genome editing in a strain that expresses the BSL1-VF protein from an unlinked locus (Extended Data Fig. 2). After crossing to the wild type, meiotic progeny carrying only the *bsl1-ko* allele without the BSL1-VF rescue construct could not be recovered (>16 recombinant tetrads tested), consistent with an essential function for BSL1.

**Fig. 1:**
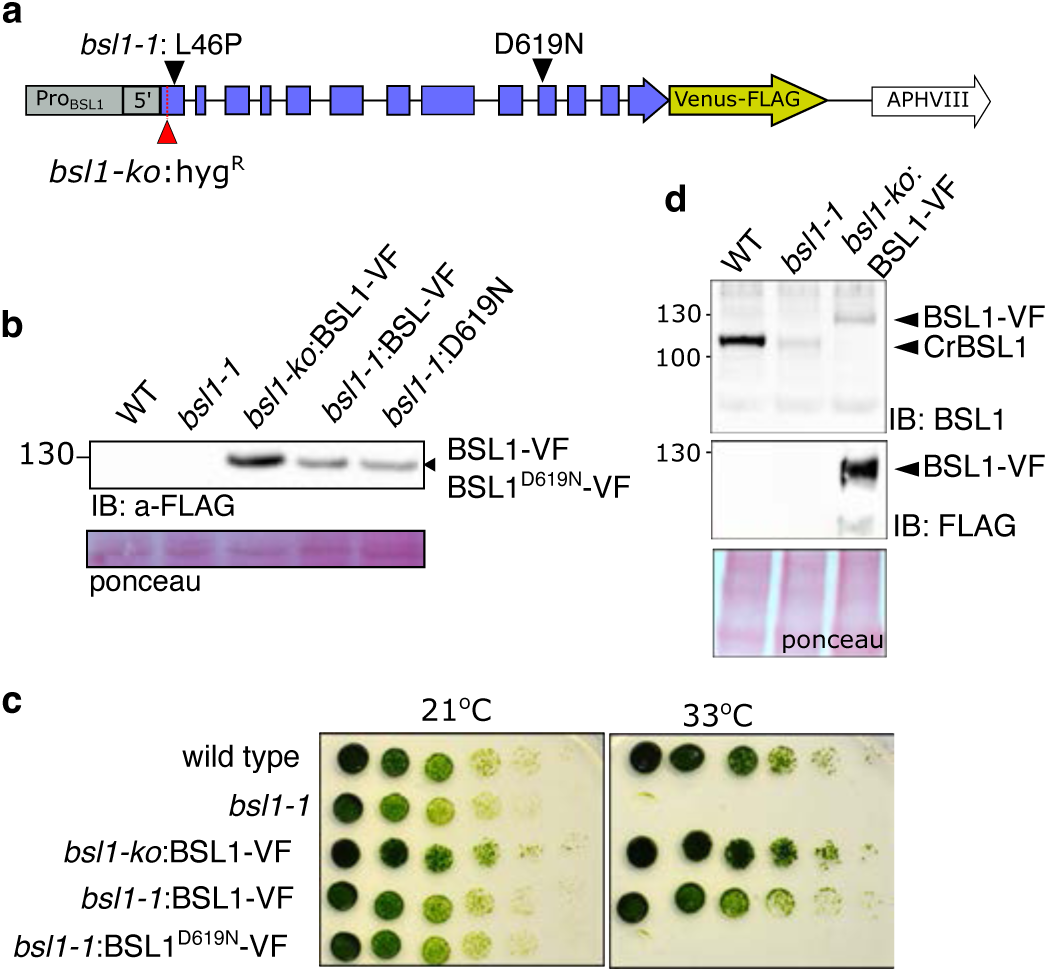
BSL1 phosphatase activity is essential for viability in Chlamydomonas. **a**, Diagram of the ProBSL1:BSL1-Venus:paroR construct shows the locations of the *bsl1-1* mutation, the insertion in *bsl1-ko*, and the D619N mutation. **b**, Anti-FLAG immunoblot shows BSL1-VF and BSL1D619N-VF in the *bsl1-1* or *bsl1-ko* backgrounds. **c**, Growth assay of the indicated strains. Cells grown at 21°C were spotted in 5-fold serial dilutions and then grown at 21°C or 33°C. Note the lethality of *bsl1-1* and of *bsl1-1:*BSL1^D619N^-VF at 33°C. **d**, Anti-FLAG and anti-BSL1 immunoblots show expression of BSL1, BSL1^L46P^-VF and BSL1-VF, and absence of endogenous BSL1 in *bsl1-ko*.

We tracked cell division in wild-type and *bsl1-1* cells at restrictive temperatures by time-lapse microscopy. The cells were synchronized in G1 at 21°C, incubated at 33°C until the onset of mitosis, and then imaged with a temperature-controlled microscope. We used a strain that expresses the microtubule end-binding protein (EB1) fused to the red fluorescent protein mScarlet (EB1-mSc), which marks cytoplasmic microtubules, the mitotic spindle, and the basal bodies^35,36^. The cells also express a bleomycin resistance protein fused to GFP (ble-GFP), which was previously shown to localize in the nucleus, except for a brief period in mitosis coinciding with spindle formation and loss of nuclear integrity^37^.

The cell cycle in wild-type Chlamydomonas is characterized by a long G1 phase, during which the cells can grow to >8-fold their starting size without DNA synthesis or cell division^22^. An interphase cell maintains a pair of cilia anchored to the cell via two basal bodies. In wild-type cells grown at 33°C, the onset of cell division was marked by the movement of the nucleus^38^ towards the basal bodies at the anterior end of the cell (Fig. 2a, Supplementary Video 1). In our movies, this nuclear movement was rapidly followed by the splitting of the two basal bodies, which moved to opposite sides of the nucleus near the future spindle poles. The ble-GFP signal disappeared from the nucleus during metaphase, indicating the breakdown of the nuclear envelope integrity. Subsequently, the cell performed nuclear division, cleavage furrow formation, and cytokinesis.

**Fig. 2:**
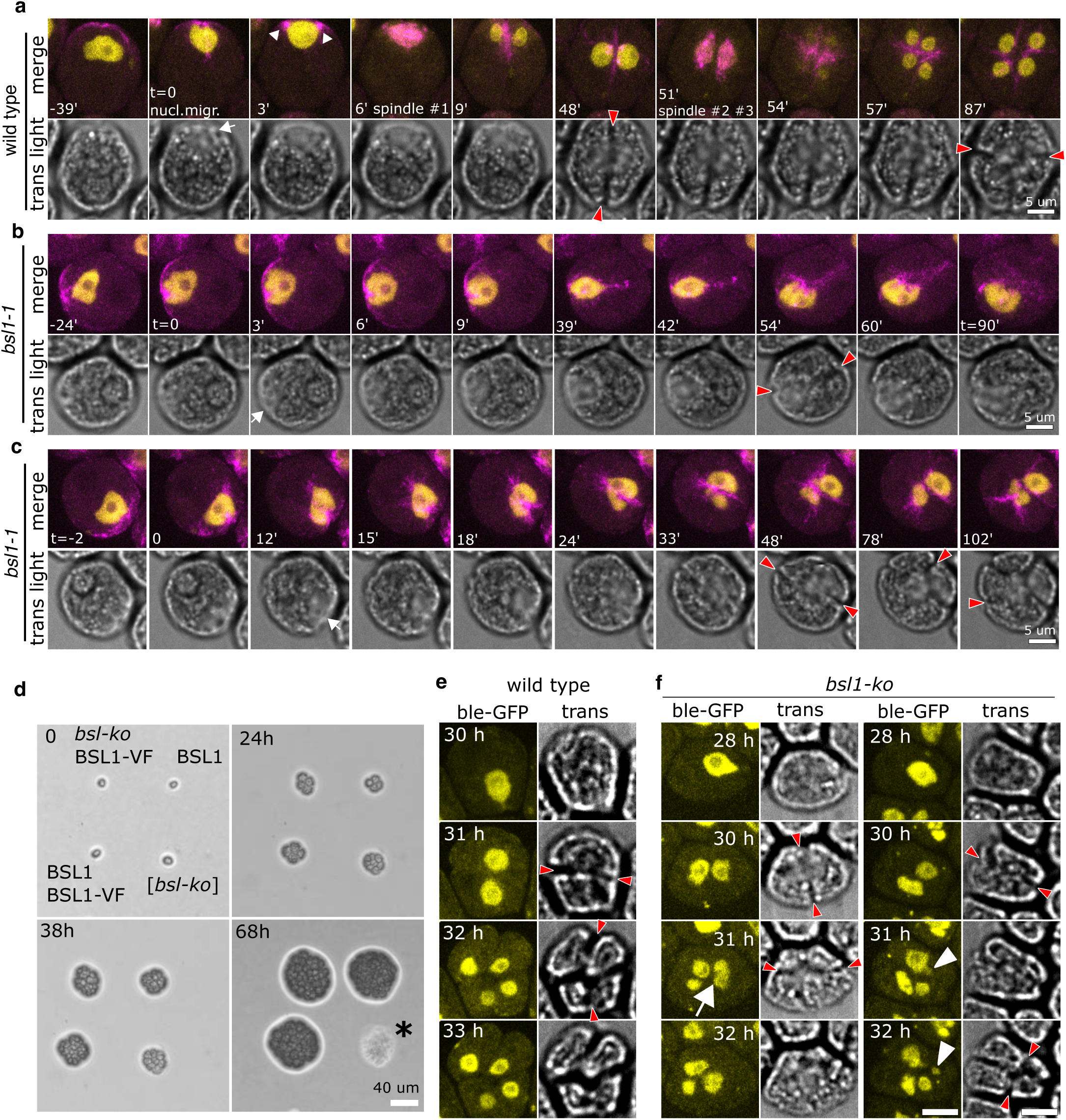
BSL1 is required for nuclear division. **a-c,** Time-lapse images of a wild-type cell (a) and two bsl1-1 cells (b and c) grown at 33°C. In each panel, the upper row shows an overlay of ble-GFP (yellow) and EB1-mScarlet (magenta), and the lower row shows corresponding brightfield images. Time (minutes) of each frame is relative to the moment the nucleus contacts basal body (t=0). **W**hite arrow: the anterior notch formed after nuclear movement. White triangles: basal body separation. Red arrowheads: cleavage furrows. Scale bar: 5 μm. **d,** Tetrad from a cross of *bsl1-ko BSL1-VF* x wild type were grown for 0, 24, 38, and 68 hrs at 22°C. Genotypes are indicated. The *bsl1-ko* genotype is inferred. The *bsl1-ko* cell performs 3-4 divisions before lysing at the 8-16 cell stage (between 38-68h, *, n=8 tetrads). Scale: 40 µm. **e-f,** Time-lapse microscopy showing ble-GFP (yellow, nucleus) in meiotic progeny from a cross (ble-GFP EB1-mSc x *bsl1-ko* BSL1-VF) at indicated hours after the start of imaging. **e,** In wild-type cells, each nuclear division results in two uniform-sized nuclei. **f,** Nuclear division failure (white arrows) and nuclear fragmentation (white arrowheads) in *bsl1-ko* cells.

The *bsl1-1* mutant grown at 33°C showed normal initial movement of the nucleus toward the basal body region (Fig. 2b, c, Supplementary Video 2). However, after nuclear migration, we observed no spindle formation, nor disappearance of ble-GFP from the nucleus. The *bsl1-1* cells did form cleavage furrows associated with EB1-mSc after a variable delay (Fig. 2b, c), suggesting that cleavage furrow formation does not require BSL1 or the residual BSL1 activity in *bsl1-1* at 33°C is sufficient for cleavage furrow formation. In most *bsl1-1* cells, the undivided nucleus was found on one side of the ingressing furrow (Fig. 2b; Supplementary Video 2). In about 20% cells, the furrow bisected the nucleus, generating nuclear fragments of varying size (Fig. 2c; Supplementary Video 3). In cells grown at 30°C, spindles formed but were randomly oriented, the duration of metaphase was significantly prolonged, and the centrin-mCherry marker, which always localizes near the spindle poles in the wild type, was clearly mis-localized away from the spindle poles in *bsl1-1* (Extended Data Fig. 3a). At the permissive temperature (21°C), most *bsl1-1* cells behaved like wild type, but we occasionally observed misoriented spindles and slightly prolonged metaphase duration (Extended Data Fig. 3c-d). These results suggest that BSL1 is essential for spindle formation, and that sufficient BSL1 activity is required for the precise organization of the spindle apparatus.

Cells carrying the *bsl1-ko* allele are inviable. To analyze the *bsl1-ko* mutant, we first crossed the complemented strain *bsl1-ko/BSL-VF* to the wild-type and monitored colony growth of meiotic progenies. From complete tetrads, where all genotypes could be assigned, we found that *bsl1-ko* progeny died at the 8-16 cell stage (Fig. 2d). We then crossed *bsl1-ko/BSL1-VF* to a wild-type strain expressing the markers ble-GFP and EB1-mScarlet and imaged meiotic progeny immediately after tetrad dissection by time-lapse microscopy (Supplementary Video 4). Under these conditions, the wild type remained viable for the duration of the movie (48h) and performed up to four rounds of cell division. Using ble-GFP as a nuclear marker, we observed tight coordination between nuclear division and cell division in the wild type, where each division resulted in two equal-sized daughter nuclei (Fig. 2e, Supplementary Video 5). In *bsl1-ko* cells, the first two nuclear divisions were indistinguishable from the wild type, probably because of parentally loaded BSL1 protein or mRNA. When *bsl1-ko* cells reached the 8-cell stage, we observed aberrant nuclear morphology and failure in nuclear division that resulted in fragmented ble-GFP signal (Fig. 2f, Supplementary Video 4, 6). Thus, the two loss-of-function mutants, *bsl1-ko* and *bsl1-1,* both display defective nuclear division.

### BSL1 localizes to segregating basal bodies and mitotic spindle poles

To understand the dynamics of BSL1 during cell division, we imaged BSL1 (BSL1-VF) together with fluorescently tagged EB1 (EB1-mScarlet) (Fig. 3a, Supplementary Video 7). Interphase cells contained BSL1-VF near the basal body region, with weak signals around the nuclear periphery. When the cells entered prophase, the BSL1-VF signal increased dramatically near the segregating basal bodies and peaked near the spindle poles in metaphase when the spindle was fully formed. In cytokinesis, BSL1-VF re-localized to a band along the cleavage furrow (Fig. 3a). In daughter cells, BSL1-VF returned to its interphase configuration, with accumulation near the basal body region and the nucleus (Supplementary Video 8).

**Fig. 3:**
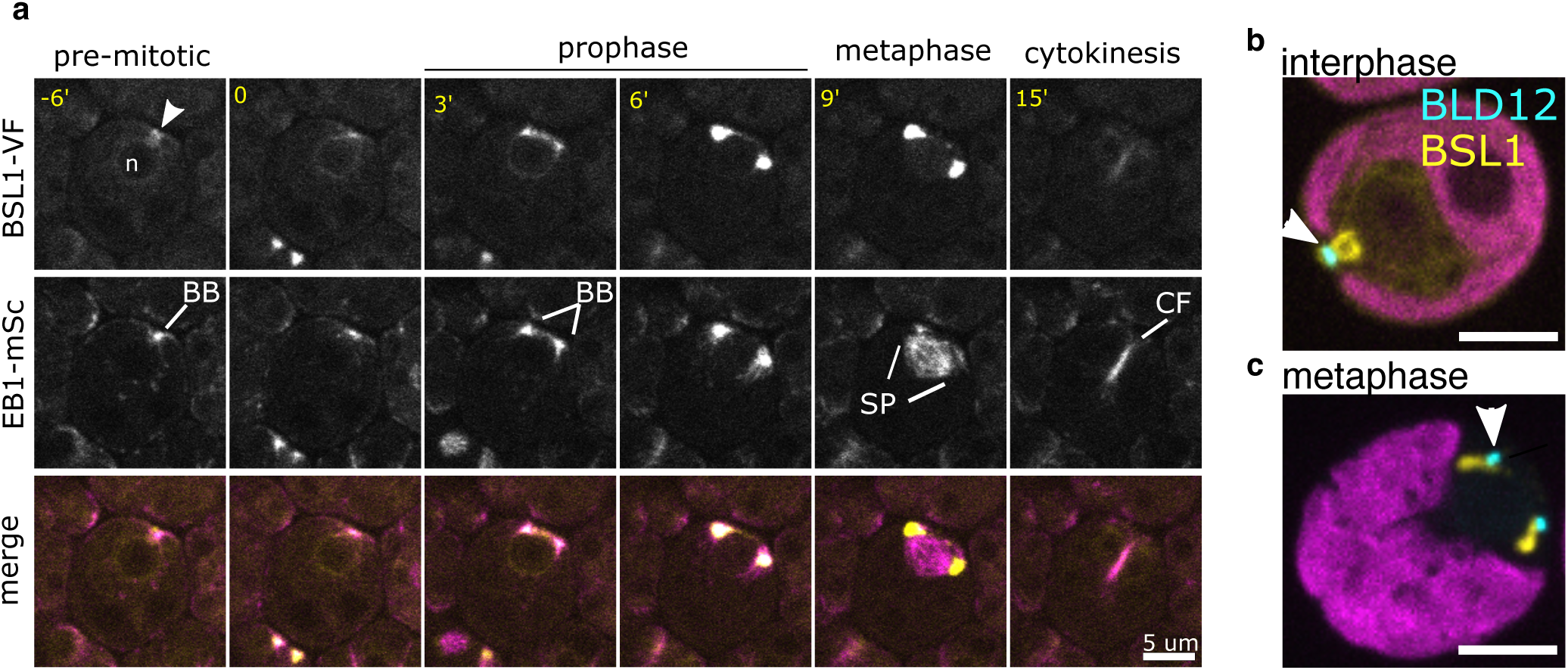
Cell cycle-dependent localization of BSL1. **a,** Confocal images of BSL1-VF and EB1-mSc at various cell cycle stages (minutes relative to nucleus contacting the basal bodies, BB). EB1-mSc labels basal bodies (prophase), spindle microtubules (metaphase), and the cleavage furrow (cytokinesis). BSL1 co-localizes with EB1 in prophase, localizes strongly near the spindle poles in metaphase, and weakly to the cleavage furrow. **b-c**, Confocal images show BLS1-mCherry (yellow), BLD12-Venus (cyan, white arrowhead), and chloroplast (magenta) in cells at interphase **(b)** and metaphase **(c)**. Scale bar: 5 μm.

In *Chlamydomonas,* the basal bodies remain attached to the cell surface during mitosis, while the mitotic spindle assembles inside the nucleus^39^. To determine the spatial relationship of BSL1 to the interphase and mitotic basal bodies, we generated a fusion between the core basal body protein BLD12 and Venus (BLD12-V). In interphase, we observed a specific signal from BLD12-V near the flagellar base in non-dividing cells (Fig. 3b), consistent with incorporation of the fusion protein into the centriole cartwheel structure^40^. In these cells, BSL1-mCh formed two filaments bridging the space between the basal bodies and the nucleus. In metaphase cells, the BLD12-V signal split in two dots, which remained in close proximity with the cell wall (Fig. 3c), and BSL1-mCh extended from the membrane-localized basal bodies towards the presumed spindle poles (Fig. 3c). The dynamic localization of BSL1 throughout the cell cycle, particularly the strong accumulation near the spindle poles, suggests that BSL1 plays a direct role in mitosis.

### The *bsl1-1* mutant is sensitive to a reduction in CDKB1 activity

Chlamydomonas contains a single CDKB gene (CDKB1), and one cyclin B gene (CYCB1) that is likely the essential activator of CDKB1 activity^17^. Inactivation of CYCB1 or CDKB1 leads to a similar mitotic arrest phenotype as in the *bsl1-1* mutant with replicated DNA, but no spindle formation and no nuclear division^16,24^. If BSL1 and CDKB1/CYCB1 act in a common pathway in mitosis, simultaneous partial loss of both genes may lead to a synthetic phenotype. We tested this possibility by crossing *bsl1-1* to the temperature-sensitive *cycb1-5* and *cdkb1-1* mutants, which are not viable at restrictive temperature (33°C)^16^, and analyzed the viability in the double mutant at the semi-restrictive temperatures (Fig. 4). To analyze the *bsl1-1 cycb1-5* and *bsl1-1 cdkb1-1* double mutants, we dissected tetrads at a permissive temperature (21°C), where all single mutants are viable without strong growth defects. After dissection, one set of tetrads was shifted to 26°C and the other set was maintained at 21°C. One colony from each tetrad was unable to grow at 26°C, suggesting that the double mutant lost viability while the single mutants were viable at the semi-restrictive temperature (Fig. 4a). The clones of a tetrad from a *bsl1-1 x cdkb1-1* cross were genotyped and tested at a lower temperature of 24°C, and the growth was scored daily (Fig. 4b,c). At 21°Cand 24°C, both single mutants were fully viable, although *bsl1-1* grew more slowly than the wild type at 21°C. The *bsl1-1 cdkb1-1* double mutant was indistinguishable from the *bsl1-1* single mutant at 21°C, but showed a strong synthetic phenotype with reduced growth and viability at 24°C, indicating a requirement for wild-type CDKB1 to maintain viability in the *bsl1-1* mutant. These genetic interactions are indicative of synthetic lethality, which suggests that BSL1 and CYCB1/CDKB1 act in concert to promote mitosis.

**Fig. 4:**
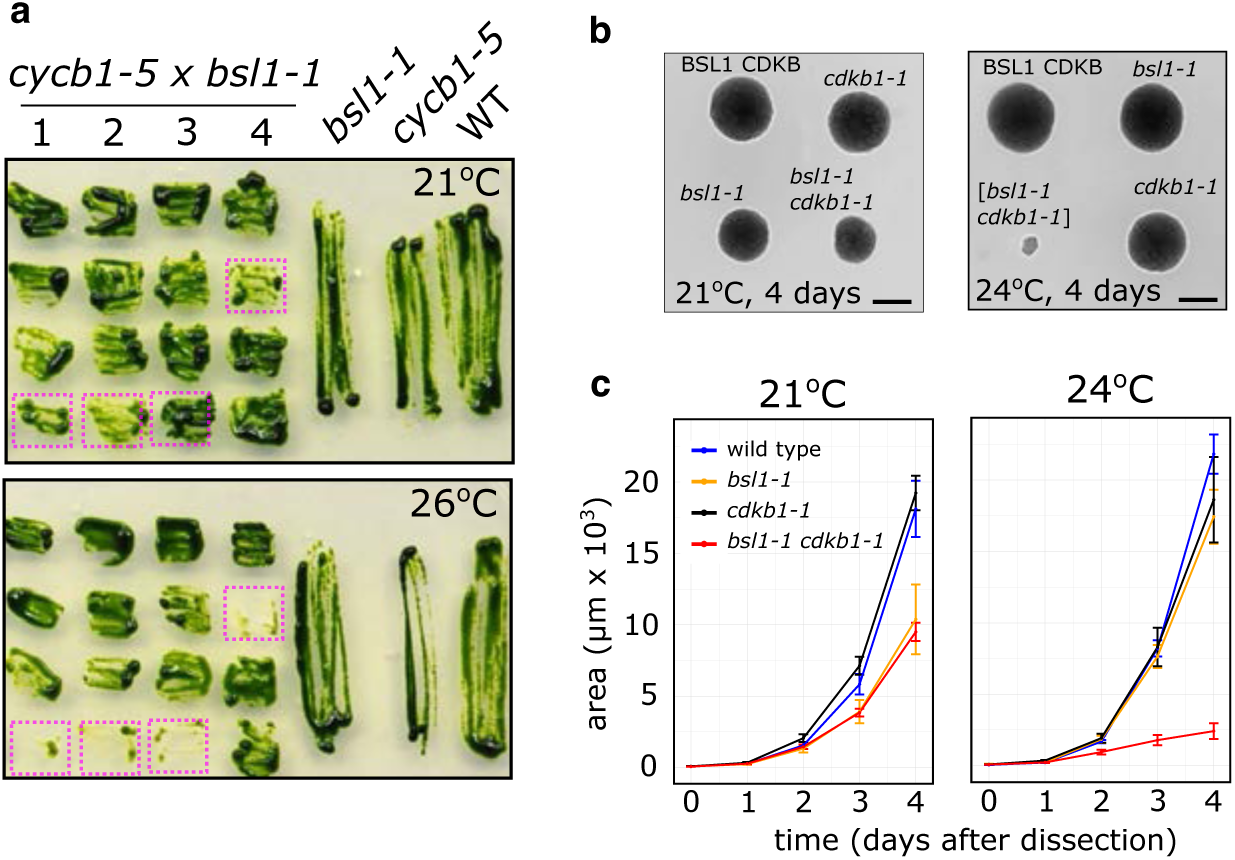
*bsl1-1* interacts genetically with *cdkb1-1* and *cycb1-5*. **a,** Tetrads from a *bsl1-1* x *cycb1-5* cross were dissected at 21°C and re-streaked to duplicate plates at 21°C and 26°C. The *bsl1-1 cycb1-5* double mutants (magenta boxes) show a synthetic lethal phenotype at 26°C. **b,** Representative images of tetrads from a *bsl1-1* x *cdkb1-1* cross grown at 21°C and 24°C for 4 days. Scale bar = 100 µm. **c,** Colony size of the indicated strains dissected from *bsl1-1* x *cdkb1-1* crosses and grown at 21°C (n=4) or 24°C (n=11). Error bars: standard error of the mean.

### BSL1 is phosphorylated in dividing cells

To test whether BSL1 is differentially phosphorylated in G1 and S/M phases, we performed anti-FLAG immunoprecipitation from synchronized BSL1-VF *cdc20-1* cells in mid-G1 (0% M phase cells) and during cdc20-induced arrest (∼30% M phase cells), with *cdc20-1* as control, and analyzed the precipitants by LC-MS/MS. The peptide coverage of BSL1 was similar in G1 (∼85%) and S/M (∼90%) cells. We identified 18 phosphorylated serine or threonine residues in BSL1 immunoprecipitated from the S/M-enriched samples (Extended Data Fig. 4a). The phosphosites were clustered either near the N-terminal Kelch domain, or in the middle region between the Kelch and the phosphatase domains. Only 3 of these sites were also detected in G1 samples, indicating increased BSL1 phosphorylation as cells enter mitosis. Nine of the S/M-specific phosphosites are conserved in Arabidopsis BSLs; four residues are also present in apicomplexan homologs (Extended Data Fig. 4b). Overall, 7 of the 18 phosphosites contained a S/T followed by a proline, which matches the minimal consensus motif of proline-directed kinases, including CDKs.

### BSL1 dephosphorylates CDKB1 pT14/pY15

To identify BSL1-dependent dephosphorylation events, we performed a global quantitative phosphoproteomic analysis in cell cycle-synchronized wild-type and *bsl1-1* cells at four time points corresponding to newborn cells (t = 0, 21°C), mid-G1 (t=6h, 33°C), early-S/M (t = 8h, 33°C) and mid-S/M (t = 10h, 33°C) in the cell cycle. Microscopic analyses indicated that the *bsl1-1* cells are delayed in cell division events (Extended Data Fig. 5a-b). Mass spectrometry analysis of flow-through peptide fractions showed accumulation of S phase (e.g. MCM3, RIR1 and CDKA) and mitotic (CYCB1, CDKB1) proteins at the 8h and 10h time points (Extended Data Fig. 5c), consistent with the accumulation of cells with S/M phase morphology at these times. Interestingly, the accumulation of CYCB1 and CDKB1 levels was delayed in *bsl1-1*. Mass spectrometry analysis of the phosphopeptide fractions indicated that hundreds of proteins showed increased or decreased phosphorylation in *bsl1-1*. These include CDKB1, which was hyperphosphorylated at T14 and Y15 (IGEG<u>pTpY</u>GK) in *bsl1-1* at the 8h and 10h time points (Extended Data Fig. 6a,b), in contrast to the decreased CDKB1 protein level at 8h (Extended Data Fig. 5c). We further quantified the intensity ratio of IGEG<u>pTpY</u>GK between the *bsl1-1* mutant and the wild type by PRM (parallel reaction monitoring) mass spectrometry^41^ (Fig. 5a, Extended Data Fig. 6c-f). The intensity of the doubly phosphorylated peptide (IGEG<u>pTpY</u>GK) was 30 to 60 times higher in *bsl1-1* compared to the wild type (Fig. 5b, Extended Data Fig. 7a-b). The same peptide phosphorylated only on Y15 was moderately increased (∼2-fold) in *bsl1-1* at the 10h timepoint (Fig. 5b). In contrast, the abundance of phosphopeptides of CYCB1, PEPC2 and CDPKK was similar between *bsl1-1* and the wild type (Fig. 5b, Extended Data Fig. 7c-e). These results indicate that BSL1 is required for efficient dephosphorylation of CDKB1 at T14 and Y15.

**Fig. 5:**
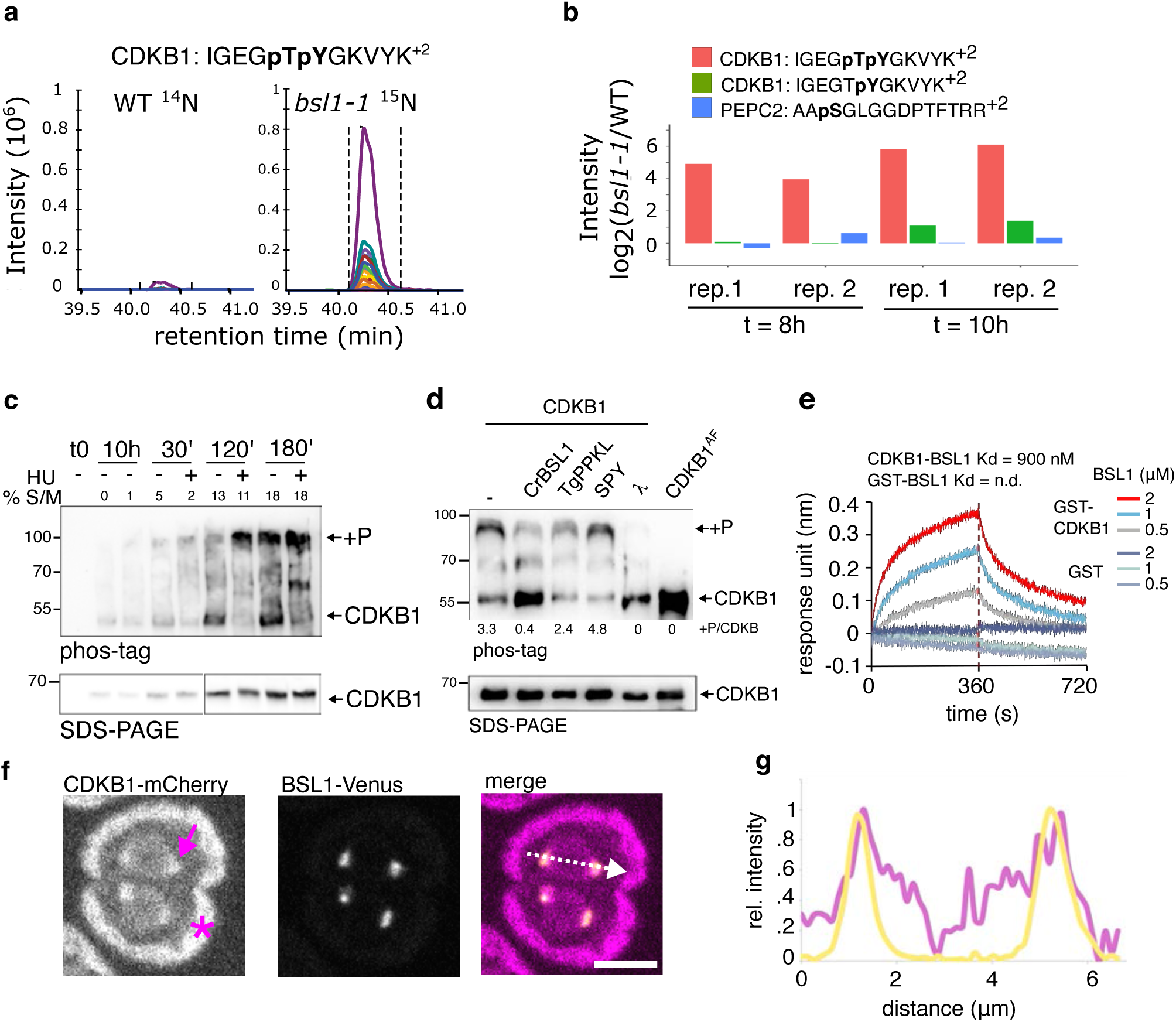
BSL1 dephosphorylates CDKB1 at T14/Y15. **a,** Quantification of CDKB1 phosphorylation in *bsl1-1* and wild-type cells after 10h at 33°C. The *bsl1-1* cells were labeled with heavy nitrogen (^15^N) and mixed with wild-type cells grown in ^14^N media before LC-MS/MS analysis. Each trace represents the intensity of a fragment ion of the IGEG**pTpY**GKVYK^+2^ peptide across the elution profile of the peptide. **b,** Quantification of the intensity ratio log_2_(*bsl1-1*/WT) of the doubly (pT14, pY15) and singly (pY15) phosphorylated CDKB1 peptides, and the control peptide from the PEPC2 protein, at the 8h and 10h timepoints. Each timepoint contains two repeats with isotope labels inversed. **c,** Parallel cultures of synchronized CDKB1-mCh cells. At 10h one culture was treated with 8 mM HU. Total protein extracts were analyzed by anti-mCh immunoblots of phos-tag and SDS-PAGE gels. +P: phosphorylated CDKB1-mCh. **d,** Wild-type CDKB1 and unphosphorylatable CDKB1^AF^ were immunoprecipitated from HU-treated cells. CDKB1-mCh was treated with recombinant phosphatase domains from Chlamydomonas BSL1 (CrBSL1), Toxoplasma PPKL (TgPPKL), or Lambda phosphatase (l). Negative controls were untreated (-) or treated with the non-phosphatase protein SPY. Dephosphorylation was measured as the intensity ratio of phosphorylated to unphosphorylated forms (+P/CDKB). **e,** Bio-layer interferometry (BLI) showing direct interaction between recombinant GST-CDKB1 and BSL1 proteins. GST-CDKB1 protein was loaded onto biosensors and dipped into indicated concentrations of purified BSL1 protein for association (time 0) and transferred into buffer (red dotted line) for dissociation. The dissociation constant (K_d_) is calculated via the ratio of the off and on rates (K_off_/K_on_). **f,** Dual-color imaging CDKB1-mCherry and BSL1-Venus in a cell with two metaphase spindles. CDKB1-mCh localizes to the spindle pole (magenta arrow). Chlorophyll autofluorescence (magenta asterisk) is captured in the mCherry channel. **g,** Normalized fluorescence intensity along a line bisecting both spindle poles (white dotted arrow in f).

In yeast and animals, phosphorylation of CDK on T14/Y15 is induced by DNA replication stress^42^. Hydroxyurea (HU), a ribonucleotide reductase inhibitor, induces replication stress in plants and animals, and was previously shown to block DNA replication in *Chlamydomonas*^43^. We found that HU increased the proportion of phosphorylated CDKB1 with little effect on the CDKB1 protein level as cells entered the S/M phase (Fig. 5c). Phosphorylation of CDKB1 was lost in cells expressing a version of CDKB1 that carries unphosphorylatable T14A and Y15A mutations (CDKB1^AF^-mCh, Fig. 5d). We performed quantitative proximity labeling with BSL1 fused to a version of BioID (BSL1-BioIDG3) that was engineered to function in the *Chlamydomonas* cytoplasm (see Methods). The result showed increased labeling of CYCB1 (Extended Data Fig. 8), which is known to form a complex with CDKB^17^. To test if BSL1 can dephosphorylate CDKB1 *in vitro*, we immunoprecipitated CDKB1-mCh from HU-treated cells as a substrate for *in vitro* phosphatase assays. The results showed that CDKB1 was dephosphorylated effectively by BSL1 and Lambda phosphatase, less effectively by *Toxoplasma* PPKL, and was not dephosphorylated after incubation with buffer only or with SPY protein as negative controls (Fig. 5d). Bio-layer interferometry assays, using recombinant proteins purified from *E. coli*, indicated that BSL1 interacts with CDKB1 *in vitro* with a K_d_ of 900 nM (Fig. 5e). Confocal microscopy showed that BSL1 and CDKB1 colocalize transiently near the spindle poles in metaphase (Fig. 5f-g), consistent with previous observation of CYCB1 localization to the spindle poles^17^. Together, the *in vivo* and *in vitro* evidence indicate that BSL1 dephosphorylates CDKB1 pT14 and pY15.

### CDKB1 phosphorylation is essential for the response to DNA replication stress

Phosphorylation at T14/Y15 is known to inhibit Cdk1/Cdc2 activity in yeast and animals^44^. To test the effect of CDKB1 phosphorylation in *Chlamydomonas*, we analyzed phosphomimetic (D14/Y15, T14/E15 and D14/E15) and unphosphorylatable (A14/F15, CDKB1^AF^) variants fused to mCherry. None of the phosphomimetic variants rescued the *cdkb1-1* mutant at the restrictive temperature (Extended Data Fig. 9a), suggesting that phosphorylation of CDKB1 blocks its function. The unphosphorylatable variant, CDKB1^AF^-mCh, complemented the *cdkb1-1* mutant but supported slower growth in colony size compared to CDKB1-mCh at restrictive temperature (Extended Data Fig. 9b), indicating that CDKB1 phosphorylation is dispensable for viability but required for optimal growth. At permissive temperature, we observed similar growth rates between the *cdkb1-1* CDKB1-mCh and *cdkb1-1* CDKB1^AF^-mCh strains, indicating that the functional CDKB1-1 is dominant over CDKB1^AF^ (Extended Data Fig. 9c). Expression of CDKB1^AF^ did not suppress *bsl1-1* (Extended Data Fig. 9d), implying that BSL1 plays additional roles besides dephosphorylating CDKB1 pT14/pY15. However, microscopic analysis indicated that CDKB1^AF^ advanced cleavage furrow formation in *bsl1-1* (Extended Data Fig. 9e), suggesting that phosphorylation of CDKB1 contributes to delayed furrowing in *bsl1-1* mutant cells. Similar to wild-type CDKB1 (Extended Data Fig. 5c), accumulation of CDKB1^AF^ was reduced in *bsl1-1* cells (Extended Data Fig. 9f), suggesting that BSL1 promotes the stability of CDKB1 independently of dephosphorylation at T14/Y15. We expressed a phosphatase-dead mutant BSL1 in the *CDKB1^AF^ bsl1-1* strain and tested the growth under a restrictive temperature. The result shows that BSL1 phosphatase activity is required even in the presence of unphosphorylated CDKB (Extended Data Fig. 9g),

Checkpoint-mediated phosphorylation of Cdk1 T14/Y15 delays the onset of mitosis until DNA replication is finished, when dephosphorylation by Cdc25 trigger exit from the delay in fission yeast^42^ and animals^45^. Since CDKB1 is the main driver of mitosis in *Chlamydomonas*, we tested the role of CDKB1 phosphorylation in response to DNA damage and replication stress. We found that *cdkb1-1* mutant cells complemented with CDKB1^AF^-mCh were highly sensitive both to HU (Fig. 6a) and to the polymerase inhibitor aphidicholin, but not to the DNA-damage-inducing drug zeocin^46^ (Extended Data Fig. 10a). The *bsl1-1* mutant cells also showed increased sensitivity to HU (Extended Data Fig. 10b), likely due to the additive effects of *bsl1-1* and HU on CDKB1 phosphorylation; this is consistent with the synthetic phenotype of *bsl1-1 cdkb1-1* double mutant (Fig. 4), supporting the role of BSL1 in exit from HU-induced mitotic arrest. We imaged *cdkb1-1* mutant cells complemented with CDKB1 or CDKB1^AF^ by time-lapse microscopy in the presence of HU (Figure 6 b-c). After 12 h incubation on control media (TAP) at 33°C, *cdkb1-1* mutant cells expressing wild-type CDKB1 or CDKB1^AF^ formed microcolonies of 8-16 daughter cells, whereas the *cdkb1-1* mutant cells did not divide after 12 h and lysed by 30 h (Fig. 6b). A low concentration (1 mM) of HU caused post-S/M lethality in CDKB1^AF^ cells without obvious impact on the wild type by 30 h, whereas a high concentration of HU (10 mM) delayed the appearance of cleavage furrows and caused lethality in all strains by 30 h (Fig. 6b-c). These results show that HU delays cell cycle progression in wild type and this delay was largely bypassed in CDKB1^AF^ cells, indicating that phosphorylation of CDKB1 T14/Y15 delays mitotic entry and cytokinesis under replication stress to prevent cell division with incompletely replicated DNA. Taken together, these results demonstrate that replication stress increases CDKB1 T14/Y15 phosphorylation, which inactivates CDKB1 and delays mitotic entry and cytokinesis; BSL1-mediated dephosphorylation activates CDKB1 and promotes mitotic entry in *Chlamydomonas*.

**Figure 6.**
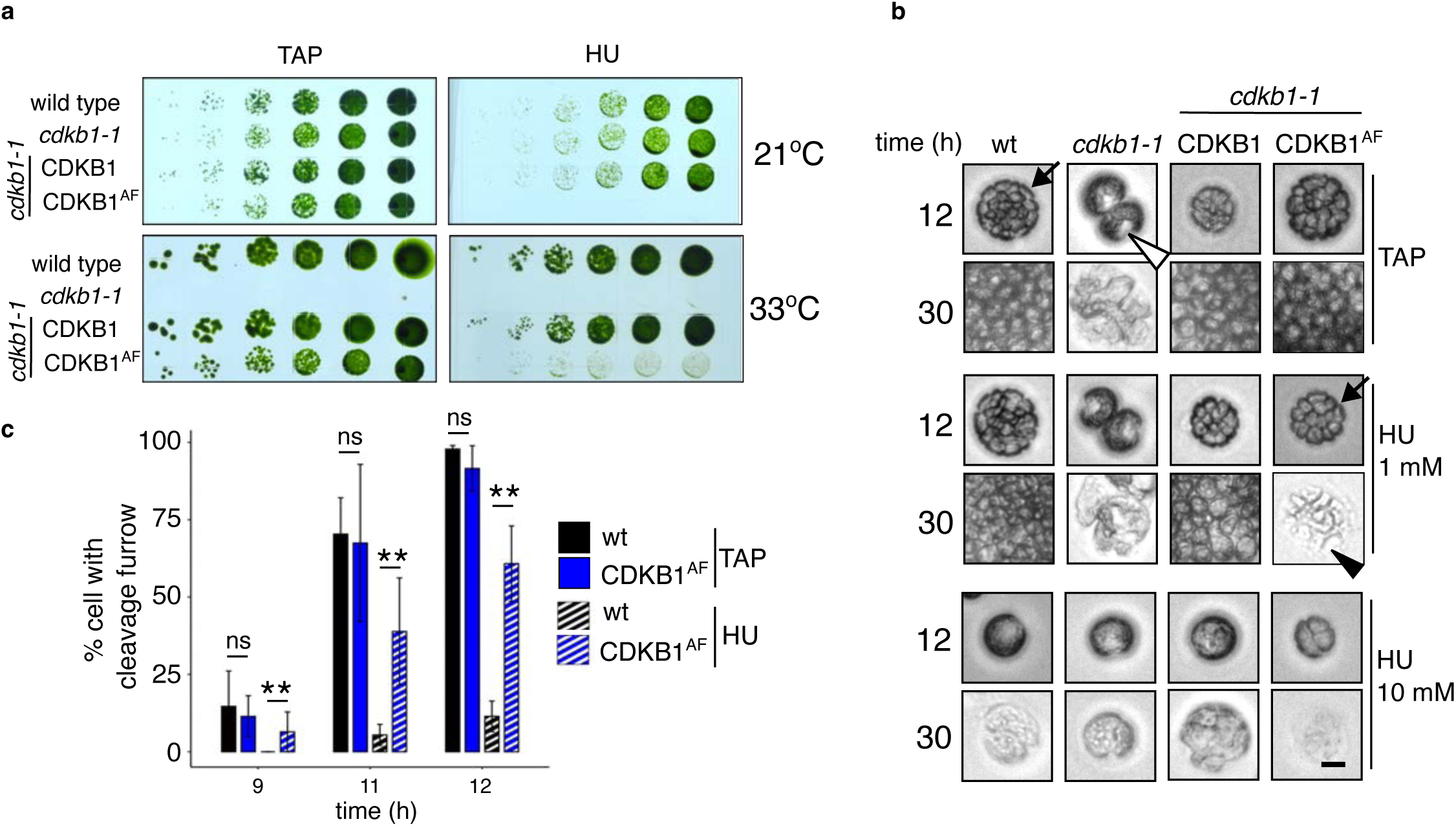
Phosphorylation of CDKB1 mediates the DNA replication stress response. **a**, Growth and survival of wild type, *cdkb1-1* and *cdkb1-1* complemented with CDKB1-mCh and CDKB1^AF^-mCh. Cells were spotted (5-fold dilution) on TAP and TAP + 1 mM HU, and grown at 22°C and 33°C. **b-c**, Microscopic analysis of cell cycle dynamics of individual cells. **b**, Selected time-lapse images of representative cells taken at 12 h and 30 h on TAP and TAP + HU media at 33°C. Black arrows point to the microcolonies containing 8-16 cells at 12 h. The *cdkb1-1* mutant arrests with a “notch” morphology (white triangle), indicating mitotic arrest. The black triangle points to representative cells undergone complete lysis. Scale: 5 μm. **c**, Quantification of cleavage furrow formation in TAP and 10 mM HU. Error bars: s.e.m. of three experiments. **: p < 0.01, chi-square test of difference between proportions.

## Discussion

Control of mitotic entry is crucial both for maintaining genome integrity and for endoreduplication in development. The mechanism of mitotic entry is well understood in opisthokonts, which include fungi and metazoans. The core mechanism involves keeping CDK inactive through phosphorylation of T14 and Y15 residues before DNA replication is complete and DNA damage is repaired, and rapidly activating CDK through dephosphorylation by CDC25. However, the mechanism controlling mitotic entry is not understood in other clades of eukaryotes^1,2^. In particular, the absence of CDC25 has created a mystery about how cell cycle progression is regulated in plants and Apicomplexa. Our study resolves this mystery by identifying BSL1 as a phosphatase that dephosphorylates and activates CDKB1, the main kinase driving mitosis in *Chlamydomonas*. We show that the inactivation of CDK by phosphorylation of T14 and Y15 is conserved, but different phosphatases have evolved to activate CDK in opisthokont and the green lineage of eukaryotes (Extended Data Fig. 11). Furthermore, unlike CDC25 and Cdk1 in mammals, which act on both G1/S and G2/M transitions^47–49^, the plant-specific BSL1-CDKB1 pathway appears to be dedicated specifically to the control of mitosis.

Our genetic, cell biological, and biochemical evidence support a model where CDKB1 and BSL1 act synergistically to drive mitosis. The temperature-sensitive alleles of *bsl1-1* and *cdkb1-1* mutants both arrest in mitosis after DNA replication and show synthetic lethal interactions at semi-restrictive conditions, suggesting that they function in a common pathway. Loss of BSL1 increases phosphorylation of CDKB1 T14 and Y15 and decreases the protein levels of CDKB1 and CYCB1, indicating that BSL1 is required for CDKB1 dephosphorylation and activation. Several lines of evidence show that BSL1 directly dephosphorylates CDKB1, including *in vitro* binding and dephosphorylation assays, colocalization of both proteins to the spindle poles during mitosis, and *in vivo* association between BSL1 and CYCB1 detected by BSL1-TurboID proximity labeling.

Phosphorylation of T14 and Y15 in CDK-family kinases interferes with catalysis and is a key mechanism for controlling cell cycle progression in fungi and animals^50^. However, a functional role for CDK T14/Y15 phosphorylation has been elusive in the plant kingdom. Mutations of T14/Y15 of CDKA1 did not affect plant response to replication stress in Arabidopsis^5^. This observation, along with the absence of CDC25, prompted the hypothesis that T14/Y15 phosphorylation is not critical for mitotic regulation in plants^5^. Here, we demonstrate that phosphorylation of T14/Y15 in CDKB1 is a key mechanism of mitotic regulation in Chlamydomonas. Inhibiting DNA replication by HU enhances the phosphorylation of CDKB1 T14/Y15, which apparently inhibits CDKB1 function, as CDKB1 containing mutations of T14 and Y15 to phosphomimetic residues was unable to rescue the *cdkb1-1* mutant. CDKB1 is essential for mitosis and cytokinesis^16^. It was reported previously that HU blocks mitosis but not cytokinesis in Chlamydomonas^43^. However, our microscopic analysis showed that HU delays cytokinesis. Consistent with HU-induced phosphorylation inhibiting CDKB1, cells expressing non-phosphorylatable CDKB1^AF^ showed a reduced delay of cytokinesis and consequently reduced survival when treated with DNA replication inhibitors. In the absence of replication inhibitors, these cells are viable, but their colonies grow more slowly than wild type, likely because some cells are lost due to premature mitosis/cytokinesis before completing DNA replication. DNA synthesis could be slowed by nutrient limitation near the end of each cell division cycle. These results indicate that replication stress causes CDKB1 T14/Y15 phosphorylation, which inhibits CDKB1 function and delays mitosis and cytokinesis till DNA replication is complete.

In fungi and animals, CDK and CDC25 form a positive feedback loop where they activate each other, and the active CDK then promotes mitotic entry. In fission yeast, the only essential function of CDC25 is to dephosphorylate CDK T14/Y15, as CDC25 is no longer essential when an unphosphorylatable Cdc2^Y15F^ is present^9^. Our study shows that the role of CDC25, i.e. dephosphorylation of pT14/pY15 of mitotic CDK, is played by BSL1 in plants. However, the mitotic arrest of *bsl1-1* mutant is not suppressed by CDKB1^AF^, suggesting that BSL1 has other essential functions in addition to dephosphorylating CDKB1 pT14/pY15. This additional function of BSL1 seems to be related to CDKB1 accumulation, as the *bsl1-1* mutation not only increased CDKB1 T14Y15 phosphorylation but also reduced the accumulation of CDKB1, CYCB1, and CDKB^AF^ proteins. BSL1 may stabilize CYCB1-CDKB1 through a phosphatase-independent mechanism, such as protein-protein interaction. However, expressing a phosphatase-dead mutant BSL1 in the *bsl1-1* CDKB1^AF^ background did not rescue viability, suggesting that BSL1’s phosphatase activity is essential even when CDKB1 is unphosphorylated at T14/Y15. Another possibility is that BSL1 dephosphorylates a protein involved in regulating CDKB1 and CYCB1 protein levels. Such regulation of both CDKB1 activity and accumulation would provide tighter control of CDKB1 function and mitotic entry.

BSL1 may have additional functions beyond regulating CDKB1. The dynamic subcellular localization of BSL1 suggests its multiple actions at different subcellular locations and cell cycle phases. The various defects in spindle organization and orientation in *bsl1-1* cells grown at semi-restrictive temperatures also suggest that BSL1 is required for both entry and proper execution of mitotic progression. Phosphoproteomic experiments showed *bsl1-1* increased phosphorylation of many proteins. As such, our genetic, microscopic, and proteomic studies suggest that BSL1 has broad functions in addition to dephosphorylating CDKB1 T14/Y15. Future studies of additional BSL1 substrates will elucidate additional signaling pathways and mechanisms that regulate cell division in *Chlamydomonas*, including the mechanism of regulating CDKB1 and CYCB1 accumulation.

BSL1 and CDKB1 appear to play different roles in cleavage furrow formation (cytokinesis), which is abolished in the *cdkb1-1* mutant but only delayed in the *bsl1-1* mutant. As noted in previous studies using HU to inhibit DNA replication^43^ or with cell cycle-blocked mutants^22^, cleavage furrow initiation does not depend on completion of DNA replication or mitosis in *Chlamydomonas*. Our analyses show that HU delays cleavage furrow formation in wild-type cells, but the delay is much reduced in cells expressing the CDKB1^AF^ protein, consistent with HU causing CDKB1 T14/Y15 phosphorylation and inactivating CDKB1. Similar to HU-treated cells, the *bsl1-1* mutant cells show a delayed cleavage furrow, which is partially suppressed by CDKB1^AF^. Together, our results support the model that DNA replication stress causes phosphorylation of CDKB1 T14/Y15 to prevent premature mitotic entry and cytokinesis before DNA is fully replicated; the BSL1-mediated dephosphorylation and stabilization of CDKB1 triggers progression through mitosis and cytokinesis.

Interestingly, CDKB1^AF^ cells showed no increased sensitivity to zeocin, which induces double-stranded DNA breaks, possibly due to distinct response pathways for different types of DNA stress. In *Chlamydomonas*, BSL1 and CDKB1 are required only for mitotic entry, whereas CDKA1 plays a dominant role in the G1/S transition^16^. Such distinct functions of CDKA and CDKB1 could enable different checkpoints to specifically control distinct cell cycle transitions. Consistent with this notion, phosphorylation of CDKA1 T14/Y15 was shown to inactivate CDKA but play no obvious role in replication stress in Arabidopsis^5^. These observations raise the possibility of a CDKB1-independent pathway for the DNA damage response and a CDKB1-dependent replication stress pathway in plants.

In animals, replication checkpoint signaling leads to CDK phosphorylation by the Wee1 kinase to safeguard DNA replication and mitotic entry^51^. WEE1 is conserved in plants. The Chlamydomonas WEE1 is expressed in S/M cells, but its function has not been reported^52^. The *Arabidopsis thaliana* WEE1 (AtWEE1) is activated by replication stress, and the loss-of-function mutants are hypersensitive to replication-inhibitory drug HU in growth but insensitive in mitotic arrest, indicating that AtWEE1 plays a role in replication stress^53^. Intriguingly, AtWEE1 was shown to mediate HU-induced phosphorylation of CDKA but not CDKB^53^. However, mutation of CDKA T14 and Y15 to non-phosphorylatable residues did not affect the plant’s sensitivity to replication stress^5^, whereas genetic evidence supports a key role for CDKBs in mitotic entry in algae and Arabidopsis^16,20^. Our study suggests that the possible WEE1-CDKB1 connection should be examined in *Chlamydomonas* and Arabidopsis in future studies (Extended Data Fig. 11). AtWEE1 was recently reported to phosphorylate the E3 ubiquitin ligase FBL17, leading to the accumulation of CDK inhibitors and inhibition of cell cycle progression^54^; how this inhibitory pathway is reversed to mediate checkpoint exit and mitotic entry remains unknown. Future studies are required to illustrate the full replication checkpoint pathways in algae and higher plants.

The regulation of mitotic entry is not only crucial for genome integrity but also important for development in multicellular organisms^20,55^. BSL1 and its homologs have evolved additional roles, particularly in receptor kinase signaling in land plants such as Arabidopsis. Members of the BSL family were initially identified in Arabidopsis as a component of the brassinosteroid signaling pathway^28^, acting downstream of the BRI1 receptor kinase to inactivate the GSK3-like kinase BIN2 by dephosphorylating its conserved tyrosine residue^29^. Recent studies revealed additional roles of BSLs in receptor kinase pathways that regulate immunity and stomata development, acting upstream of mitogen-activated protein kinase modules, with signaling specificity enabled by phosphocoding and scaffolding^30–32^. In Arabidopsis, CDKB1 is known to play a role in mitotic entry and endoreduplication^20^. An intriguing possibility to be tested in the future is that receptor kinase-mediated signaling may regulate endoreduplication through a BSL-CDKB1 connection in certain land plants. Our study in Chlamydomonas suggests that BSL1 is a cell cycle regulator that has evolved additional signaling functions during the evolution of land plants. It is conceivable that BSLs may act as a signaling hub that integrates diverse developmental and environmental signals to regulate cell division, differentiation, and endoreduplication in plants.

Our findings raise interesting questions about the evolution of BSL/PPKL phosphatases. The BSL/PPKL family is conserved in plants and Apicomplexa but absent in fungi and metazoans^8^, and thus their presence coincides with the absence of CDC25 in evolution. BSL/PPKL is essential for viability in *Toxoplasma gondii*, and is thus considered an ideal target for anti-parasite drug development^27^. Our *in vitro* phosphatase assay shows that TgBSL/PPKL can dephosphorylate Chlamydomonas CDKB1. The function of BSL/PPKL in mitosis seems ancestral and conserved, and such conservation of a cell cycle regulator would add a key piece of supporting evidence for the algal origin of Apicomplexa. Future studies of BSL/PPKL hold great potential for improving agricultural productivity and managing human diseases caused by Apicomplexan parasites.

## Acknowledgments

We thank Dr. Andrey Malkovskiy at the Carnegie Institution Imaging Facility. We thank Dr. Amy Ikui (Brooklyn Collage) for generously providing the *cycb1-5* mutant and Dr. Fred Cross (The Rockefeller University) for generously providing the ble-GFP strain. The CDKB-mCh (pKA1) and CDKB^AF^-mCh (pKA17) plasmids were constructed by K.C. Atkins and generously provided by Dr. Cross. Research reported in this publication was supported by funding from the European Union’s Horizon 2020 research and innovation programme (2014-2020) under the Marie Skłodowska-Curie grant agreement No 798198 (FT), the National Institute of General Medical Sciences of the National Institutes of Health (under Award Numbers R01GM066258 to ZW, R01GM135706 and S10OD030441 to SX), National Science Foundation (MCB 1818383 to MO and John R. Pringle), and the Carnegie Endowment Fund to the Carnegie Mass Spectrometry Facility.

## Author contributions

Conceptualization: FT, FA, ZW

Methodology: FT, AR, YA, YF, MO, SX, ZW

Investigation: FT, ZW

Visualization: FT, ZW

Funding acquisition: FT, FA, ZW

Project Administration: FT, FA, ZW

Supervision: ZW, AG, FA

Writing – original draft: FT, FA, ZW

Writing – review & editing: FT, SX, MO, FA, ZW

## Data availability

All data are available in the main text or the supplementary materials. Mass spectrometry raw data will be deposited at ProteomeXchange before publication.

## Supplementary Materials

## Supplementary Methods

### Strains and culture conditions

Wild type *Chlamydomonas* strains used are isogenic with CC-4402 (mt+) and CC-4403 (mt-). The *bsl1-1* mutant was isolated after UV mutagenesis^18^, and backcrossed eight times to CC-4402/4403 before analysis. The *cycb1-5* and *cdkb1-1* were described^17^. Chlamydomonas crosses were performed according to standard methods^56^ and meiotic products were dissected using a Zeiss Axioscope A1 equipped with a microdissection needle. Strains were propagated on TAP (Tris Acetate Phosphate) media at 21°C with constant illumination (∼20 µmol m^-2^ s^-1^). For selection, paromomycin (Sigma P9297) and hygromycin B (EMD Millipore 400052) were used at 10 µg/ml in TAP agar. For metabolic labeling, strains were passaged twice on TAP media containing heavy ^15^NH_4_Cl (Cambridge Isotopes, NLM4675) instead of regular ^14^NH_4_Cl before being used for experiments.

Asynchronous populations of cells in liquid culture were synchronized in G1 by washing cells once in water and once in Tris Phosphate (TP) media, followed by incubation in TP media, 12 h at 21°C in the dark. For time-course experiments, one of two protocols was used. In the plate protocol, ∼2.5*10^7^ G1-cells were spread on a 12 x 12 cm TAP agar plate and incubated at 33°C under 50 µmol m^-2^ s^-1^. In the liquid culture protocol, TAP media was inoculated to ∼OD_750_ 0.1 and grown at 33°C under 120 µmol m^-2^ s^-1^. Cell division was scored visually by the appearance of cleavage furrows by brightfield microscopy using a 40x objective. In the plate protocol, wild-type cells typically initiated division at 8-10 h, in the liquid protocol at 11-13 h.

### Plasmid construction

The BSL1 gene (Cre01.g050850) was amplified from CC-4403 genomic DNA using bsl-1F/R and bsl-2F/R primers, and introduced into pCR-Blunt II-TOPO (Thermo 450245) to generate pFT1. A 2kb promoter fragment was amplified from CC-4403 genomic DNA with bsl2kb-F and bsl2kb-R and introduced in pCR-Blunt II-TOPO to generate pFT2. The plasmid pFT4, for expression of BSL1 in *Chlamydomonas* from the synthetic H/R promoter [Pro_H/R_:gBSL1-Venus—APHVIII] was generated by amplifying the CrBSL1 from pFT1 with bsl-cp4F and bsl-cp4R and introducing into pMO449^[57]^ by Gibson assembly (New England Biolabs, E2611S). The native promoter version pFT40 (Pro_BSL1_: BSL1-mVenus-3xFLAG) was generated by inserting the BSL1 promoter from pFT2 into PfoI+HpaI digested pFT4 by Gibson assembly. The phosphatase-dead version BSL1^D619N^ was generated by circular mismatch PCR from pFT40. The CDKB1-mCh (pKA1) and CDKB1^AF^-mCh (pKA17) plasmids were generously provided by Dr. Cross (Rockefeller University). The phosphomimic CDKB1 variants (CDKB1^DY/TE/DE^-mCh) were generated by mismatch PCR from pKA1. The BioIDG3 sequence corresponds to the G3 generation in the directed evolution of TurboID from BioID (see supplementary table 1 in Branon et al. ^58^), including the mutations: Q65P, I87V, R118S, E140K, Q141R, A146Δ, S150G, L151P, V160A, T192A, M209V, I305V. The BioIDG3 sequence was codon-optimized for *Chlamydomonas* and combined with pFT40 to generate the constructs pFT57 (Pro_BSL1_:BSL1-BioIDG3-NeonGreen) and the control construct pFT58 (Pro_BSL1_: BioIDG3-NeonGreen).

### Transformation

Electroporation was performed essentially as described^59^ with 1.2 µg of linearized plasmid per approximately 2*10^8^ cells. 10 µl plasmid digestion reaction was performed in FastDigest buffer (Thermo Scientific). Cells were spun down and washed twice in electroporation buffer [10 mM N-cyclohexyl-2-aminoethanesulfonic acid, pH 9.25, 40 mM sucrose, 10 mM sorbitol]. 125 µl of cell suspension was added to the digested plasmid, mixed by pipetting, and added to a 2 mm electroporation cuvette (Bulldog Bio). Electroporation was performed using a NEPA21 instrument (Bulldog Bio) with settings: Voltage (poring pulse 250V/transfer pulse 20V), length (poring 8 msec/transfer 50 msec), interval (poring 50 msec/transfer 50 msec), # pulses (poring 2/transfer 5), decay (poring 40%/transfer 40%), polarity (poring +/transfer +/- alternating). Impedance was generally 0.33-0.37 Ohm. Electroporated cells were transferred to 8 mL TAP + 40 mL sucrose and incubated ∼20h, 22°C in dim light (<5 µmol m^-2^ sec^-1^), then spun down and plated on selective media. Transformants were selected after 7 days. Transformation with EcoRV-digested pFT40 typically resulted in ∼100 colonies after selection at 33°C and ∼1000 colonies when selected at 21°C on 10 µg/ml paromomycin.

### CRISPR-guided knock-out of CrBSL1

A culture of the complemented bsl1:BSL1-VF strain (50 ml, OD_750_=0.2) was spun down and resuspended in gametic autolysine enzyme (prepared from CC-4402/ CC-4403 gametes) and incubated 1h at 22°C. Loss of cell wall integrity was verified by treatment with 0.25% Triton X-100, which caused bursting of autolysine-treated cells. After autolysine-treatment, cells were heat-shocked for 30 min at 40°C with gentle shaking. During heat-shock treatment, RNP complex was prepared: sgRNA (100 pmol/µl, Integrated DNA Technologies) and SpCas9 (61 pmol/µl, Integrated DNA Technologies) were mixed in the following ratio: sgRNA 1.2 µl, SpCas9 1 µl, duplex buffer 2.8 µl [30 mM HEPES pH 7.5, 100 mM KOAc] and incubated 30 min, 37°C then kept on ice. 500 ng of APHVII PCR product (hygromycin resistance gene, amplified from plasmid pRAM103^60^ with MOP804 MOP626) was added to PCR tubes. Heat-shocked cells were spun down 3 min, 2000 rpm, washed twice in TAP+40 mM sucrose and resuspended in ∼500 µL TAP + 40 mM sucrose. Cell suspension (125 µl) was added to the APHVII PCR product, mixed by pipetting, and then transferred to the sgRNA/Cas9 mixture. The mixture (125 µl) was used for electroporation with the above settings. Impedance in TAP+40 mM sucrose was ∼0.15 Ohm. Electroporated cells were transferred to 8 mL TAP + 40 mL sucrose and incubated ∼20h, 22°C in dim light (<5 µmol m^-2^ sec^-1^), then spun down and plated on TAP+10 ug/ml hygromycin B. Transformants were selected after 7 days, under 50 µmol m^-2^ sec^-1^light. Eight of 12 hygromycin-selected transformants contained the APHVII gene inserted in either the native BSL1 locus or in the BSL1-VF transgene. No clone with simultaneous disruption of both loci was identified.

### Timelapse microscopy

For fluorescence timelapse microscopy, cells were pre-grown in TAP media at 21°C in constant illumination (20 µmol m^-2^ s^-1^) and synchronized in G1 by the TP-DARK method. Synchronized cells were grown in TAP liquid (∼10-11 h) or TAP agar (8-9h) before imaging. Temperature-sensitive strains were grown at 33-34°C. Agar sandwiches were prepared by melting TAP-agar (1.5%) and pipetting 275 µl on an 18 x 18 mm coverslip (Avantor) and covering with a second coverslip. When the agar had set, one coverslip was removed and 2 µl of cell suspension was spotted on the agar surface. Mutant and wild-type control cells were spotted next to each other on the same agar slab. A 24 x 60 mm coverslip (Avantor, size 1.5) was placed on top of the cells and the agar edges were sealed with grease to prevent drying of the agar during imaging. The 24 x 60 mm coverslip was placed on an inverted spinning disc confocal microscope (Olympus) equipped was a 100x silicone immersion objective (NA 1.35) and a heated stage insert (Tokai-Hit) set to hold 33-34°C inside the chamber. mVenus and GFP fluorophores were excited with a 514 nm laser using a 440/514/561 nm beamsplitter and a 550/49 nm emission filter. mCherry was excited with a 561 nm laser using a 440/514/561 nm beamsplitter and a 605/64 nm emission filter. For time-lapse imaging, images were collected every 3 minutes. This time interval ensured that every mitotic spindle would be captured, while minimizing photo-toxicity to the cells.

For brightfield timelapse microscopy, cells were synchronized as above and spotted on TAP-agar plates. For drug treatments, hydroxyurea (Sigma-Aldrich, H8627), aphidicholin (EMD Millipore, 178273), or zeocin (InvivoGen, ant-zn-1) was added to TAP agar. Cells were incubated at 33°C, ∼80 µmol photons m^-2^ s^-1^, and imaged with a 10x objective.

### Statistical analyses

Statistical tests were performed using R and plotted with the ggplot2 package^61^. All sample means comparisons (t-test, ANOVA) were two-sided. Tukey’s post-hoc tests were performed with the R package multcompView.

### Immunoprecipitation (IP)

Cells were harvested by centrifugation, transferred to 50 mL tubes and washed once in IP buffer [40 mM HEPES pH 7, 100 mM KCl, 100 sorbitol]. The wet pellet was weighed and an equal amount (w/v) IP buffer added. The resulting slurry was added dropwise to liquid nitrogen to make frozen cell pellets that were stored at -70°C. The pellets were cryomilled in a Retsch cell grinder, 3 min @ 30 Hz and. Approximately 400 mg milled powder per IP was thawed 2 min at room temperature, then transferred to ice, and 1.5 ml ice-cold IP buffer with protease inhibitors (Sigma #A32695) and phosphatase inhibitor (Pierce #32957 1 tablet/5 ml) was added. The suspension was homogenized by pipetting and clarified by centrifugation 10 min 10,000 xg, 4°C. Supernatant was added to 100 µl anti-FLAG magnetic beads (Sigma #M8823) and the samples incubated 1h, 4°C with end-over rotation. Beads were washed 4x with IP buffer. Bound proteins were eluted in SDS-PAGE loading buffer (Laemmli), without DTT [1% SDS, 10% glycerol 62.5 mM Tris-HCl pH 6.8], 10 min, 65°C. A second elution was done in 100 µl 2x Laemmli [2% SDS, 50 mM DTT], 10 min, 65°C. Generally, >90% of BSL1-FLAG eluted in the first elution. Sampled were separated on 7.5% Tris-glycine SDS-PAGE gels for in-gel digestion and mass spectrometry.

### Immunoblot

Immunoblot analysis was performed according to standard methods, using monoclonal mouse anti-FLAG antibody (Sigma F1804, clone M2), and polyclonal rabbit anti-mCherry (Abcam, ab167453) primary antibodies. The anti-BSL1 serum was raised against exon13 of CrBSL1 (amino acids 770-885 in Cre01.g050850.t2.1 Phytozome v5.6) fused at its N-terminus to maltose-binding protein. Total extracts for immunoblot analysis were prepared by mixing 30 µl of 2.5x Laemmli buffer [2.5% SDS (w/v), 125 mM Tris-HCl pH 6.8, 12.5% glycerol, 0.0125% bromophenol blue] supplemented with 50 mM DTT, with 10 mg wet cell pellet and heating 10 min at 65°C. Extracts were clarified by centrifugation and approximately 7.5 µL loaded per sample. Zn^2+^ phos-tag gel analysis was performed according to the method of Kumar^62^.

### Phosphatase assay

The phosphatase domain of *Chlamydomonas* BSL1 (amino acids 507-885 in Cre01.g050850.t2.1) and *Toxoplasma* PPKL (amino acids 532-934 in UniProt S7UP60) were fused at the N-terminus to EcoRI/NotI-digested pSUMO DUet vector to create N-terminal 6xHis and SUMO tags. 6xHis-SUMO-SPY from *Arabidopsis* (AT3G11540) was included as a negative control. Proteins were produced in *E. coli* BL21 cells and partially purified by nickel column chromatography. For *in vitro* phosphatase assays, cells expressing CDKB1-mCh were synchronized as above and incubated 13 h at 31°C 180 µmol m^-2^ s^-1^ white light. Hydroxyurea (8 mM) was added after 10 h. Cells were harvested by centrifugation, and the pellet was resuspended in 2 volumes ice-cold TBS (17.5 mM Tris-HCl pH 7.6, 137 mM NaCl), added dropwise into liquid nitrogen, and milled to a fine powder. To precipitate CDKB1-mCh, 150 mg powder was resuspended in 400 µl TBS with cOmplete protease inhibitors (Roche 04693132001, 1 tablet per 50 ml) and phosphatase inhibitors (Pierce #32957 1 tablet/5 ml), and 15 ul RFP-Trap beads (Chromotek, RTMA-20) were added. After 1 h incubation, rotating at 4°C, beads were washed 5x with 1 ml TBS, and 2x with phosphatase reaction buffer [50 mM Tris-HCl pH 6, 5 mM MnSO_4_], then aliquoted into tubes. 1 µg purified protein (BSL1-PP, PPKL-PP, SPY), or 200 U lambda phosphatase (Santa Cruz, sc-200312A) was added to CDKB1-mCh bound to RFP-Trap beads in a 20 µl volume and incubated 3 h at room temperature. The beads were washed 5x with 1 mL TBS and heated 10 min at 65°C in 30 µl 2.5x Laemmli buffer [2.5% SDS (w/v), 125 mM Tris-HCl pH 6.8, 12.5% glycerol, 0.0125% bromophenol blue]. Approximately 5 µl of the eluate was analyzed by Zn^2+^ phos-tag gel according to the method of Kumar Kumar^62^. Blots were probed with polyclonal anti-mCherry antibody (Abcam, ab167453).

### BioIDG3 proximity labeling

Strains expressing BSL1-BioIDG3-NeonGreen and the control BioIDG3-NeonGreen in the *cdc20-1* background were labeled with ^15^N and synchronized in G1 as above. 500 ml TAP or TAP-^15^N cultures (OD_750_ ∼ 0.1) were transferred to 33°C, 120 µmol light m^-2^ s^-1^ at t = 0. Biotin was was dissolved in TAP or TAP-^15^N and added to the cultures at t = 11 h to a final concentration of 250 µM. Cultures were harvested at t = 14 h by centrifugation 5 min, 4000 xg. The pellet was weighted and resuspended in 200 µl buffer (20 mM Tris-HCl pH 7.4, 150 mM NaCl) per 100 mg pellet and added dropwise to liquid nitrogen. Pellets were milled in a Retsch tissue homogenizer (M1000, 3 min, 30 Hz) to a fine powder. Sample and control powder were mixed in equal amounts by weight. Proteins were precipitated as above. Precipitated proteins were dissolved in 6M guanadinium, with sonication, reduced (5 mM tris(2-carboxyethyl)phosphine, 45 min, 55°C) and alkylated (10 mM iodoacetamide, 45 min, room temp. dark). Protein concentration was determined (BioRad 5000001) and 1 µg trypsin (Sigma, T6567) was added to 20 ug protein in 1.5 M guanidnie NH_4_HCO_3_ and incubated 12 h at 37°C. Trypsin was added again and incubated 3 h 37°C. The reaction was acidified by adding 1% formic acid and de-salted over C18 Sep-Pak resin (Waters #023590). Desalted peptides were resuspended in 1 ml PBS [137 mM NaCl, 2.7 mM KCl, 10 mM Na_2_HPO_4_, 1.8 mM KH_2_PO_4,_ pH 7]. 100 µl M280 beads (Invitrogen 11205D) was equilibrated in PBS and added to the peptides and the solution was incubated 1h at r.t. with constant agitation (800 rpm). The beads were washed 1x with PBS (5 min at r.t. 800 rpm), 1x with PBS + 5% acetonitrile (5 min at r.t. 800 rpm) and 1x with ddH_2_O + 5% acetonitrile (5 min at r.t. 800 rpm). Biotinylated peptides were eluted 3x with 200 µl 0.2% trifluoroacetic acid, 0.1% formic acid, 80% acetonitrile (each elution 5 min, 75°C, 800 rpm). Elutions were pooled and peptides were dried by evacuation and resuspended in 200 µl 5% acetonitrile + 0.1% trifluoroacetic acid and de-salted using ZipTips (MilliporeSigma ZTC18S960). De-salted peptides were dried and analyzed by LC-MS/MS.

### Bio-layer interferometry (BLI)

The BLI assay was conducted using a Gator-Plus instrument (Gator Bio) following a protocol described previously^63^. In brief, anti-GST biosensors (Gator Bio # 160042) were loaded with GST-CDKB1 or GST expressed in *E. coli*. These sensors were then immersed in wells containing varying concentrations of purified recombinant BSL1. After a 360-second association phase, the sensors were transferred to 1x PBS for dissociation. The dissociation constant (K_d_) was calculated using Gator Bio’s data analysis software, based on the K_off_/K_on_ ratio.

### Phosphopeptide enrichment

Wild type and *bsl1-1* cells were pre-grown in TAP-^14^N or TAP-^15^N agar plates and inoculated in liquid media and synchronized as above. G1 cells were plated on TAP-agar and incubated at 33°C in continuous light. Cells were harvested by scraping the plates with ice-cold water, added to crushed ice and pelleted by centrifugation in 50 ml tubes. An aliquot of harvested cells was kept for microscopic scoring of cell division. The pellet was transferred to microfuge tubes and weighed, resuspended in buffer (17.5 mM Tris-HCl pH 7.6, 137 mM NaCl with phosphatase inhibitors (Pierce #32957 1 tablet/5 ml; 200 µl buffer used per 100 mg pellet). Wild type and *bsl1-1* cells were mixed in equal amount based on pellet weight. The mixture was added dropwise to liquid nitrogen and stored at -70°C. The frozen cells were cryomilled using a Retsch tissue homogenizer (M1000, 3 min, 30 Hz). Protein extraction buffer [100 mM Tris-HCl pH 8, 2% w/v SDS, 1.2 v/v Triton X-100, 20 mM EGTA, 20 mM EDTA] was added (3 ml per gram powder), vortexed 1 min and heated 10 min, 55°C. Insoluble material was pelleted by centrifugation, 15 min 16,000 xg. The supernatant was extracted with phenol (pH 8) and the phases separated by centrifugation 15 min 16,000 xg. The organic phase was re-extracted twice with 50 mM Tris-HCl pH 8. Proteins were precipitated by adding ice-cold 100 mM ammonium acetate in MeOH with overnight incubation at -70°C. Precipitated proteins were pelleted by centrifugation, 15 min 16,000 xg. The pellet was washed twice with 100 mM ammonium acetate in MeOH, twice with MeOH and air dried 10 min to remove traces of MeOH. Proteins were resuspended in 10% SDS with 15 min at 55°C followed by 15 min in a heated water bath (42°C) sonicator (Fischer Scientific Ultrasonic Bath) and finally by sonication with a metal probe (Branson Digital Sonifier, 10% amplitude, 10 sec). This resulted in near-complete solubilization of precipitated proteins. The average yield was 2.3 mg protein per 100 mg cell pellet. The solution was made 5% SDS, 50 mM triethylammonium bicarbonate (TEAB) pH 8.5 and proteins were reduced (5 mM tris(2-carboxyethyl)phosphine, 45 min, 55°C) and alkylated (10 mM iodoacetamide, 45 min, room temp. dark). Approximately 5 mg protein was digested using S-Trap midi columns (Protifi # C02-midi-10) according to the manufacturer’s instructions, with 1:20 (w/w) trypsin (Sigma, T6567) per sample. Eluted peptides (∼1 mg per sample) were de-salted using C18 Sep-Pak resin (Waters #023590). Phosphopeptide enrichment was performed using TiO_2_ tips (Thermo #32993) according to the manufacturer’s instructions and analyzed by LC-MS/MS. Phosphopeptides enrichment was >95%. Peptide concentrations in flow-through fractions were normalized and analyzed by LC-MS/MS.

### Mass spectrometry and data analysis

Peptides were analyzed by liquid chromatography–tandem mass spectrometry (LC-MS/MS) on an Easy LC 1200 UPLC liquid chromatography system connected to an Orbitrap Eclipse quadrupole-orbitrap (Thermo Fischer) or a Q-Exactive HF hybrid quadrupole-orbitrap (Thermo Fischer) mass spectrometer in data-dependent acquisition mode (DDA). Precursor scans were between 375-1600 m/z and the 20 most intense multiply charged peptides were selected for fragmentation by higher-energy collision dissociation (HCD) with normalized collision energy (NCE) 27. Tandem mass spectrometry peak lists were extracted by an in-house PAVA script, and were searched by ProteinProspector^64^ for metabolically labeled peptides essentially as reported previously^65^, against a database constructed from *Chlamydomonas* proteins (Phytozome v5.6) with a decoy database of reversed peptides. Precursor and MS/MS tolerance was set to 10 ppm and 20 ppm, respectively. For phosphopeptide identification, Serine/Threonine/Tyrosine phosphorylation was used as variable modification. Phosphorylated CDKB1 peptides (IGEG**pTpY**GKVYK and IGEGpT**pY**GKVYK) were detected in DDA mode. To quantify the relative intensity of phosphorylated CDKB1 peptides in *bsl1-1* and wild-type cells, the peptides were re-injected with the same Easy LC 1200 UPLC liquid chromatography gradient and analyzed by the Eclipse quadrupole-orbitrap in parallel reaction monitoring (PRM) mode, as described^41^. The intensity of MS2 fragment ions across the elution peaks was plotted using the Skyline software^66^, as described previously^41^. Raw data is uploaded to proteomexchange.org with accession code xxx.

### Sequence alignments and accession numbers

Multiple sequence alignments were performed by the Muscle algorithm in Jalview (cite: doi: 10.1093/bioinformatics/btp033). Arabidopsis thaliana (Ath) AthBSL1 NP_192217.2, AthBSL2 NP_001184935.1, AthBSL3 NP_180289.3, AthBSU1 NP_171844.6. Toxoplasma gondii PPKL XP_002368432.1, Plasmodium falciperum PPKL XP_001348804.1. Phytozome.org (v.5.6) identifiers for *Chlamydomonas* genes mentioned in this study: BSL1 (Cre01.g050850), CDKB1 (Cre08.g372550), CYCB1 (Cre08.g370401), MCM3 (Cre06.g295700), RIR1 (Cre12.g492950)

---

**Supplementary Table 1.**
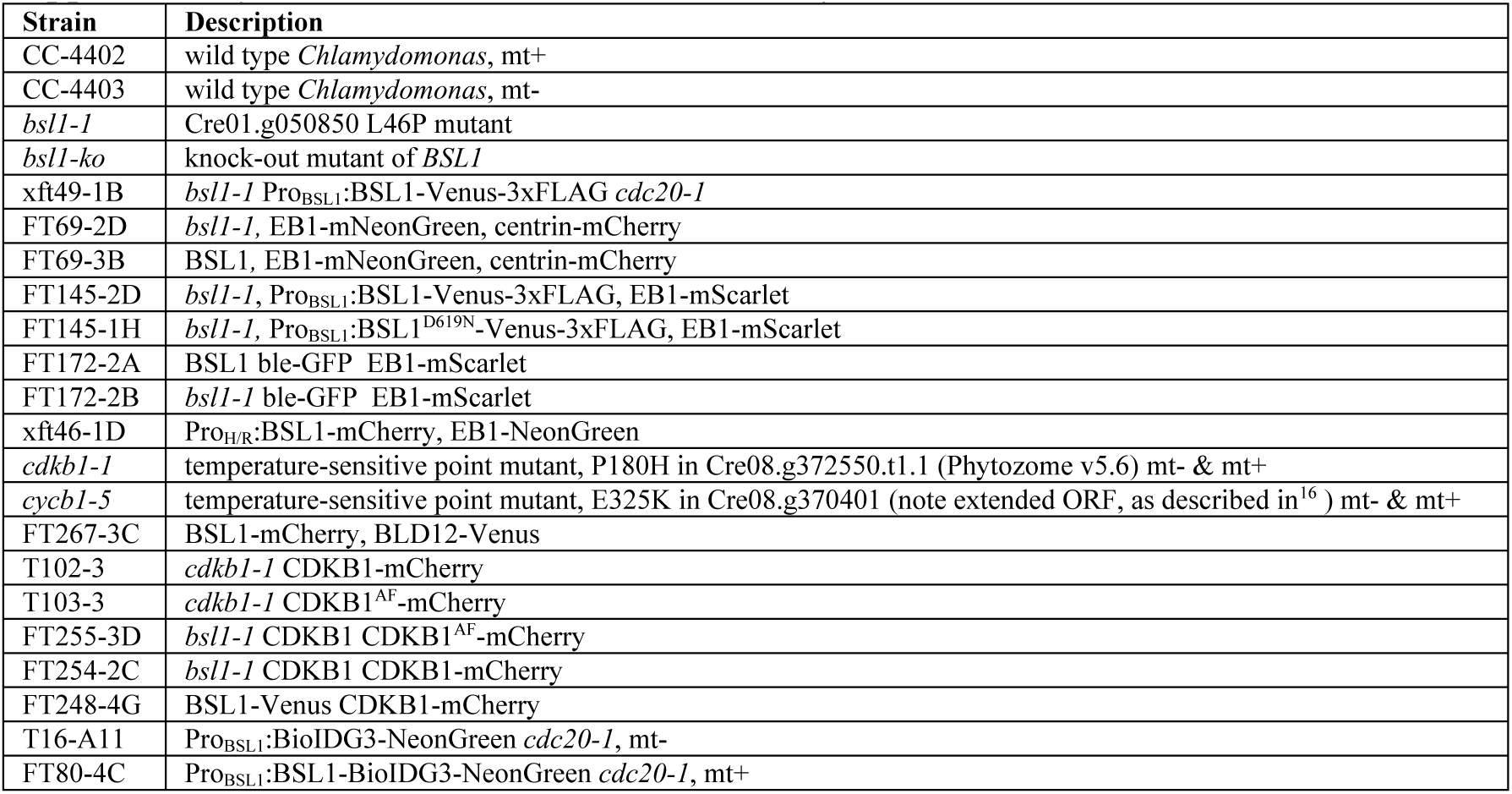
Strains used in this study.

**Supplementary Table 2.**
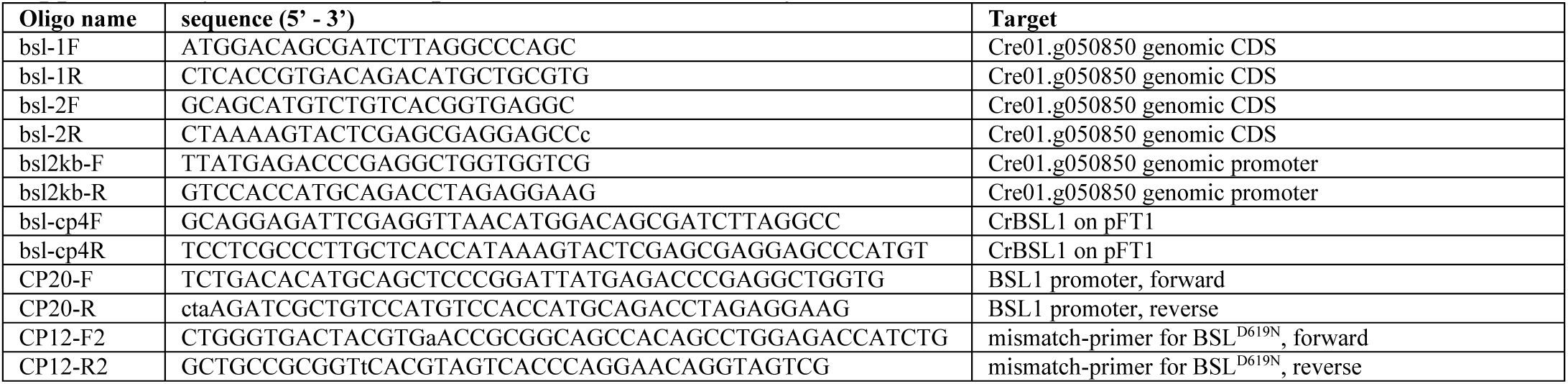
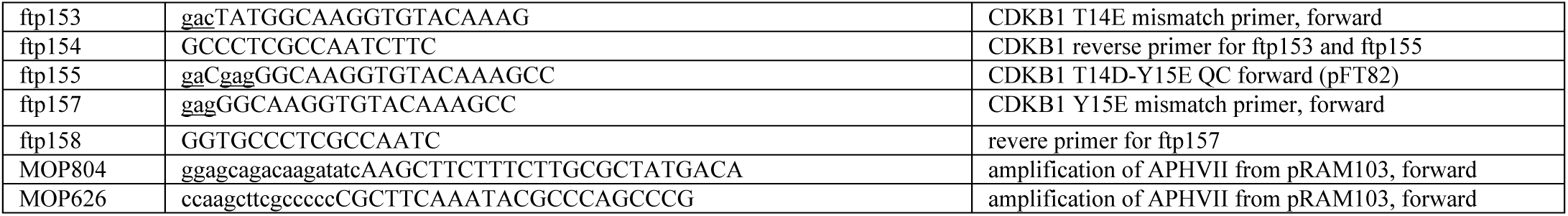
PCR primers used in this study.

## Legends for Supplementary Videos

**Supplementary Video 1. Dynamics of ble-GFP and EB1-mScarlet in mitosis.**

Wild-type cell expressing ble-GFP (a nuclear marker) and EB1-mScarlet (a microtubule marker), grown at 33°C. Acquisition every 3 min. Scale 10 um.

**Supplementary Video 2. Mitosis dynamics of ble-GFP and EB1-mScarlet in *bsl1-1*.**

A *bsl1-1* cell expressing ble-GFP and EB1-mScarlet, grown at 33°C. Acquisition every 3 min. Chloroplast division at t=36 min, cleavage furrow at t=72min. The nucleus does not divide. Scale 10 μm.

**Supplementary Video 3. Mitosis dynamics of ble-GFP and EB1-mScarlet in *bsl1-1*.**

A *bsl1-1* cell expressing ble-GFP and EB1-mScarlet, grown at 33°C. Acquisition every 3 min. Cleavage furrow at t=51min. No spindle forms. The ingressing furrow bisects the nucleus. Scale 10 μm.

**Supplementary Video 4. Mitosis dynamics of ble-GFP in *bsl1-ko*.**

One tetrad from a cross: *bsl1-ko*:BSL1-VF x EB1-mScarlet ble-GFP. The *bsl1-ko* segregant was assigned based on the lysis phenotype at the 8-16 cell stage (Fig. 3; Extended Data Fig. 3a). The presence of BSL1-VF was assigned based on the localization of BSL1-VF to spindle poles in mitotic cells. *bsl1-ko* is recessive in the presence of BSL1-VF. Genotypes of meiotic products are A) BSL1-VF EB1-mSc, B) BSL1-VF, C) ble-GFP EB1-mSc, D) *bsl-ko* ble-GFP. The second allele of *bsl1-ko* may be in either A or B. scale: 10 µm.

**Supplementary Video 5. Division of wild-type cells.**

A wild-type cell expressing ble-GFP (nuclear marker) was imaged every hour. scale: 10 µm.

**Supplementary Video 6. Division of *bls1-ko* cells.**

A *bsl1-ko* cell expressing ble-GFP was imaged every hour. scale: 10 µm.

**Supplementary Video 7. Localization of BSL1-Venus through in mitosis**

A *bsl1-1* cell complemented with BSL1-Venus (yellow), co-expressing EB1-mScarlet (magenta) undergoing mitosis. Acquisition every 3 min.

**Supplementary Video 8. Localization of BSL1-Venus through a multiple-fission cycle**

A *bsl1-1* cell complemented with BSL1-Venus (yellow). Acquisition every 1 h. The cell undergoes two divisions, producing four daughters. At exit from S/M, BSL1-Venus returns to the anterior end of the cell.

