## Extended Data Figures 1-11 for "Mitotic entry is controlled by the plant-specific phosphatase BSL1 and cyclin-dependent kinase B"

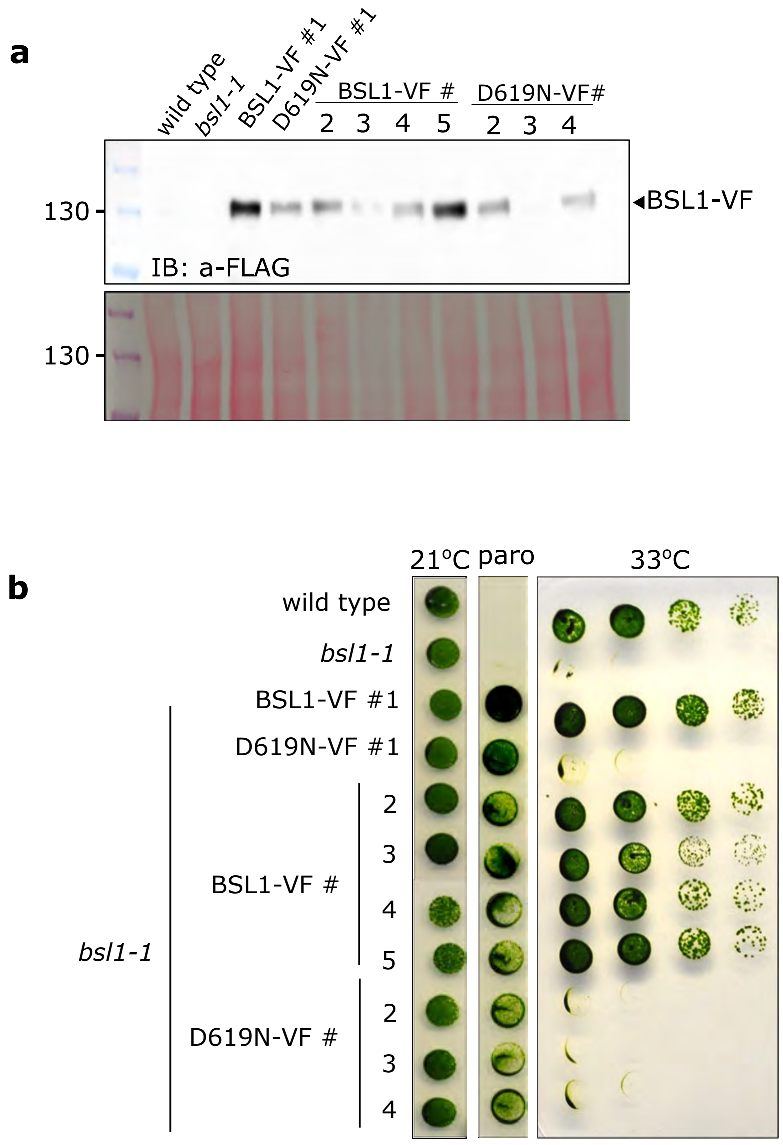


**Extended Data Fig. 1. BSL1 phosphatase activity is essential for viability in Chlamydomonas.**

**a**, Anti-Flag immunoblot analysis of five *bsl1-1*:BSL1-VF and four *bsl1-1*:BSL1$^{D619N}$-VF transformants. **b**, Growth assays at 21°C with or without paromomycin (paro) and at 33°C without paromomycin. Paromomycin is the selection marker linked to BSL1-VF and BSL1$^{D619N}$-VF.

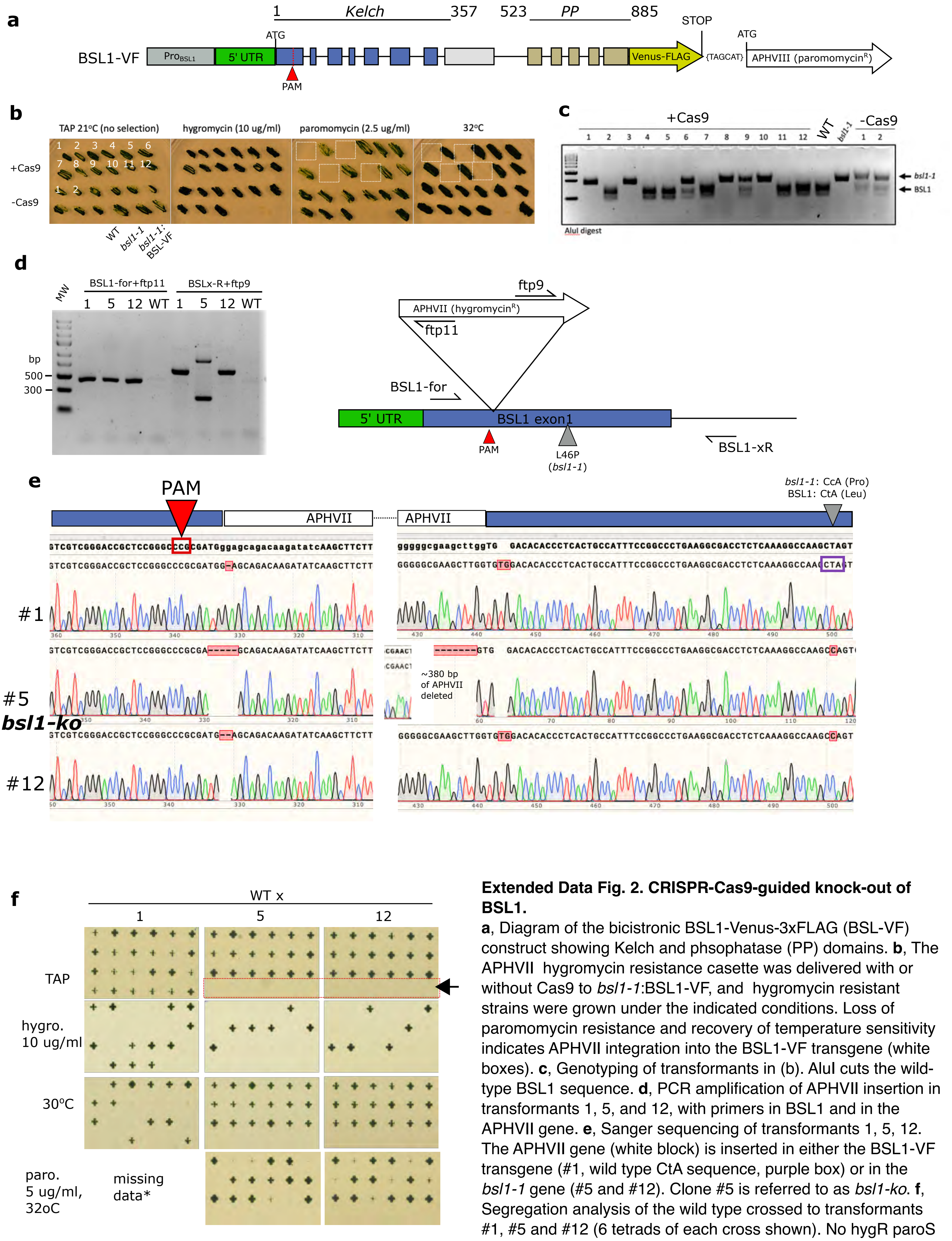


**Extended Data Fig. 2. CRISPR-Cas9-guided knock-out of BSL1.**
**a**, Diagram of the bicistronic BSL1-Venus-3xFLAG (BSL-VF) construct showing Kelch and phsophatase (PP) domains. **b**, The APHVII hygromycin resistance casette was delivered with or without Cas9 to *bsl1-1*:BSL1-VF, and hygromycin resistant strains were grown under the indicated conditions. Loss of paromomycin resistance and recovery of temperature sensitivity indicates APHVII integration into the BSL1-VF transgene (white boxes). **c**, Genotyping of transformants in (b). AluI cuts the wild-type BSL1 sequence. **d**, PCR amplification of APHVII insertion in transformants 1, 5, and 12, with primers in BSL1 and in the APHVII gene. **e**, Sanger sequencing of transformants 1, 5, 12. The APHVII gene (white block) is inserted in either the BSL1-VF transgene (#1, wild type CtA sequence, purple box) or in the *bsl1-1* gene (#5 and #12). Clone #5 is referred to as *bsl1-ko*. **f**, Segregation analysis of the wild type crossed to transformants #1, #5 and #12 (6 tetrads of each cross shown). No hygR paroS recombinants (*bsl1-ko*, red box) could be isolated in #5 and #12, indicating that disruption of the *bsl1-1* locus, without the BSL1-VF rescue construct, is lethal.

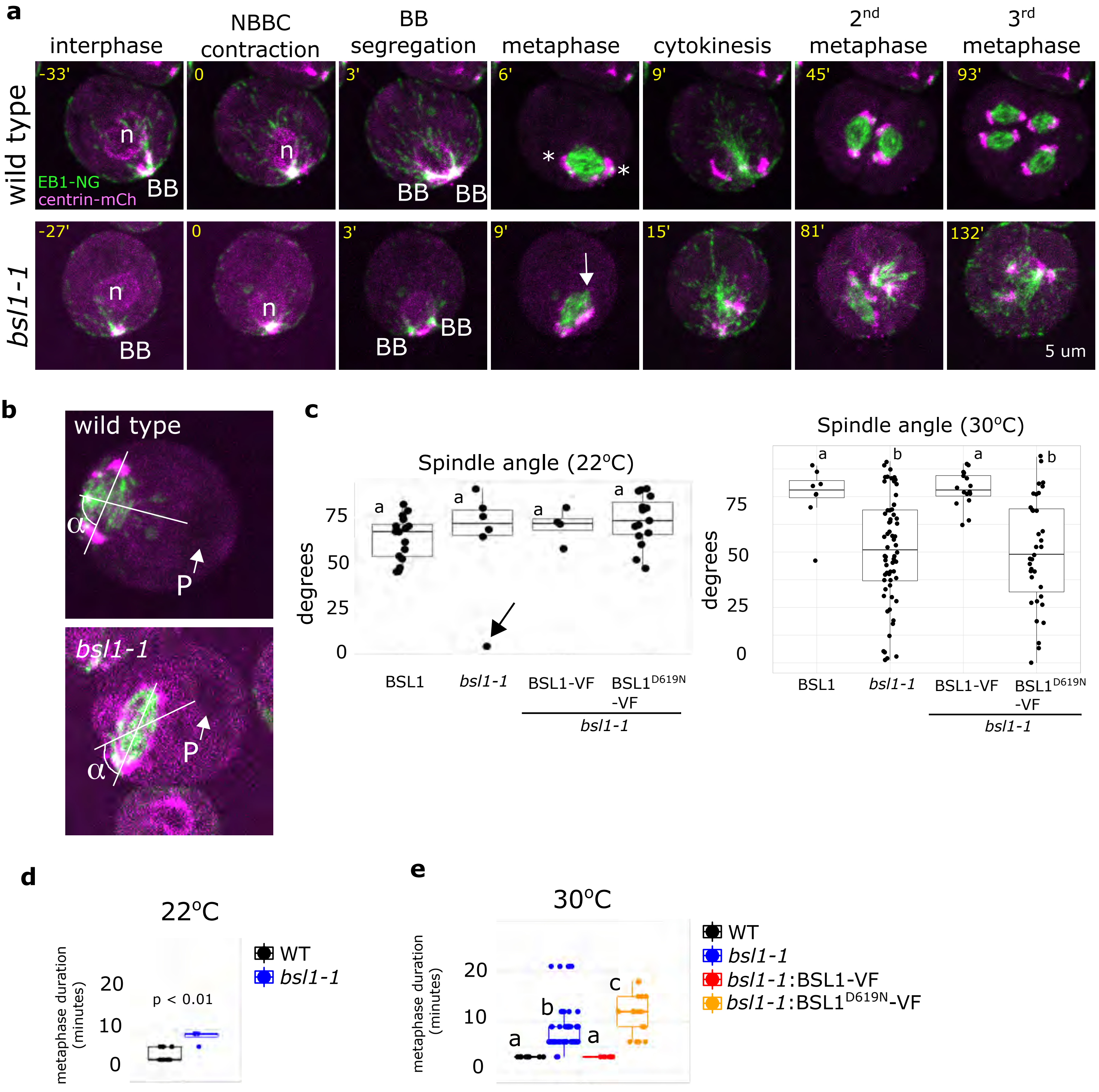


**Extended Data Fig. 3: Spindle misorientation in the *bsl1-1* mutant grown below 33°C.**
**a**, Wild type and *bsl1-1* mutant cells co-expressing EB1-mNeonGreen (green) and centrin-mCherry (magenta) imaged by time-lapse microscopy. The time (minutes) of each image is relative to the nuclear-basal body contact (NBBC, time 0). Nucleus (n) and basal body (BB) are marked. In wild type, the BBs segregate (time 3'), the mitotic spindle forms with centrin-mCh localized to the spindle poles (white asterisks). In *bsl1-1*, NBBC and BB separation appears similar to the wild type but one spindle pole does not recruit centrin-mCherry in metaphase (white arrow); the cleavage furrow MTs are more disorganized compared to the wild type; and the 2nd and 3rd divisions display highly aberrant spindles. Scale: 5 um. **b**, The orientation of the mitotic spindle was measured as the angle (α) between one line through the centers of the spindle and pyrenoid (P) and one line through the two spindle poles. **c**, Quantification of spindle orientation in cells grown at 21°C (one-way ANOVA, df=3, F=1.1) and ~30°C (one-way ANOVA, df=3, F=9.2). Occasional spindle misorientation was observed in *bsl1-1* at 21°C (black arrow). **d**, Quantification of the duration of metaphase at 21°C (two-sided t-test on sample means, p<0.01). **e**, Quantification of the duration of metaphase at 30°C (one-way ANOVA (df=3, F=12.01). c-e: Box plots represent median, 25th/75th percentile with whiskers max 1.5 * inter-quartile range. Letters indicate significance levels , Tukey post-hoc p<0.05.

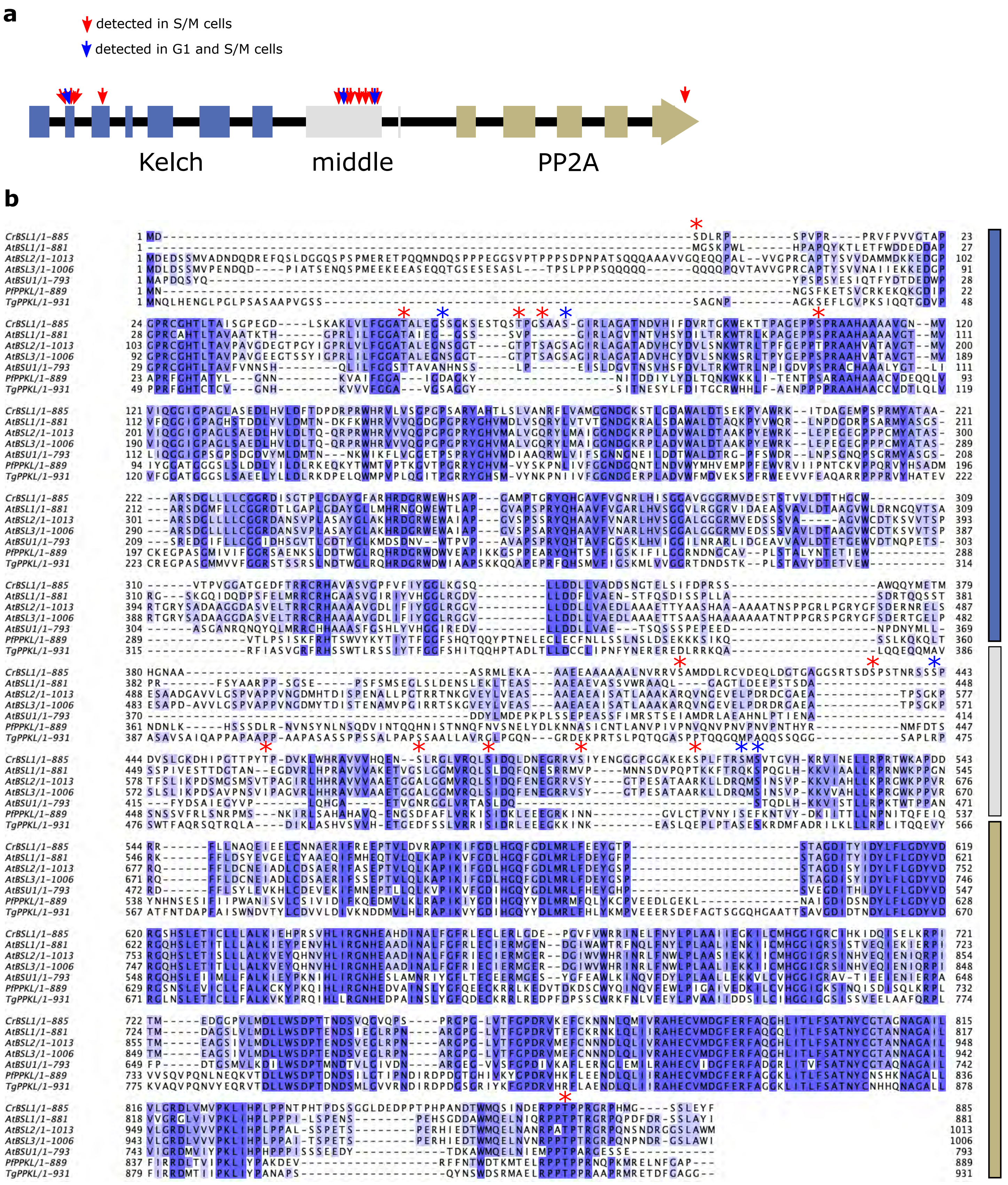


**Extended Data Fig. 4: BSL1 phosphorylation sites. a**, Diagram of BSL1 protein structure shows the relative locations of phosphosites identified only in S/M phase cells (red arrow) and those identified in both S/M and G1 cells (blue arrows). **b**, Multiple sequence alignment of *Chlamydomonas* BSL1, *Arabidopsis* BSL1,2,3 and BSU1, PPKL from *Toxoplasma gondii* (Tg) and *Plasmodium falciparum* (Pf). Asterisks: phosphosites, color codes as in (a).

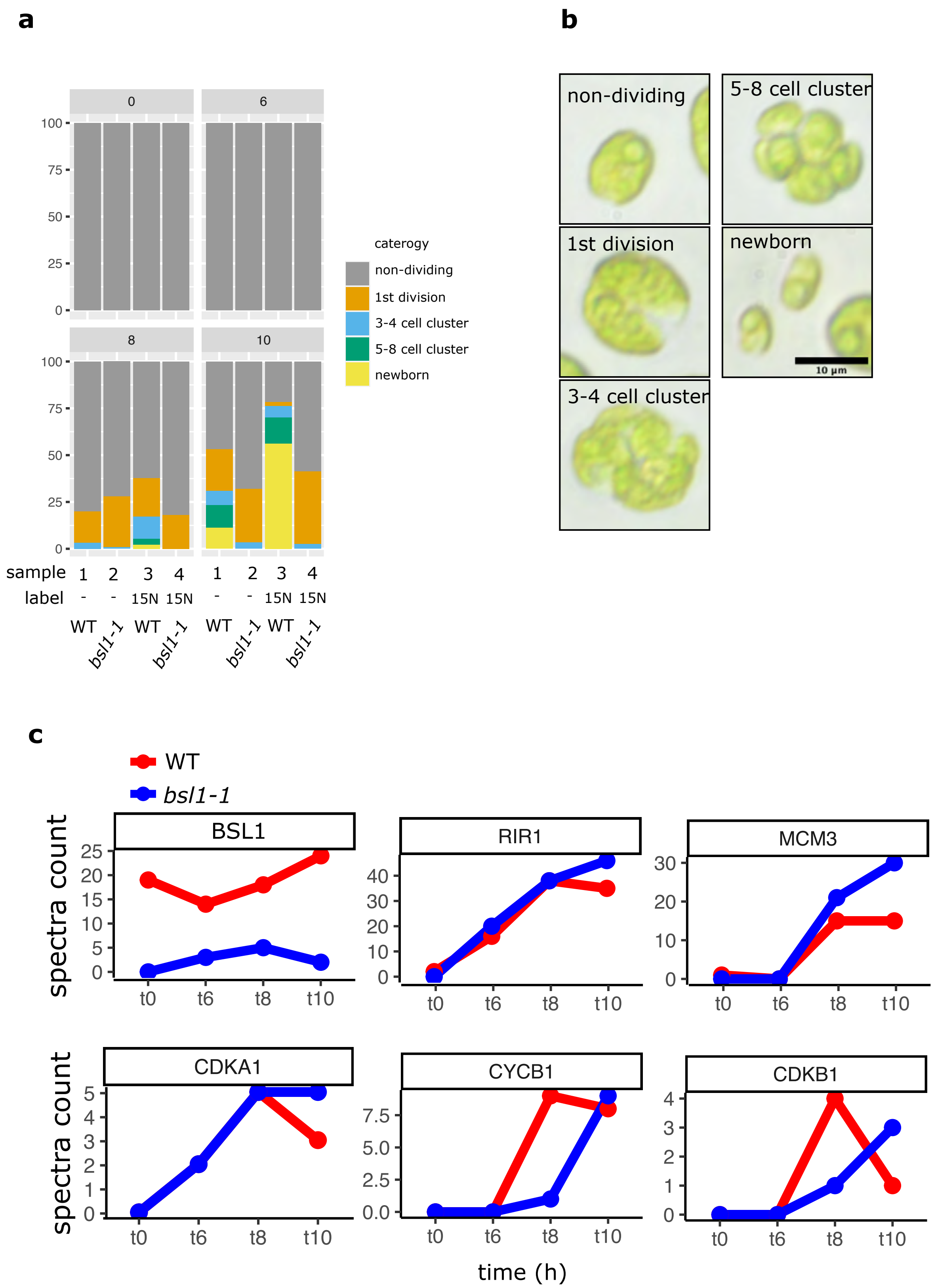


**Extended Data Fig. 5. Proteomic analyses of the *bsl1-1* mutant.**
**a**, Quantification of cell cycle progression at 0, 6, 8 and 10 hrs at the restrictive temperature. Over 100 cells were counted per timepoint per sample. For quantitative mass spectrometry analysis, the Light and heavy N samples were combined as: 1+4 (FORWARD), and 2+3 (REVERSE). **b**, Example of cells from the respective category. **c**, Spectra counts of selected proteins in wild type and *bsl1-1*. Spectra counts of BSL1$^{L46P}$ are reduced compared to wild type. Ribonucleotide reductase (RIR1) and the helicase MCM3 and CDKA accumulate with similar kinetics in wild type and *bsl1-1*. Accumulation of CYCB1 and CDKB1 is delayed in *bsl1-1*.

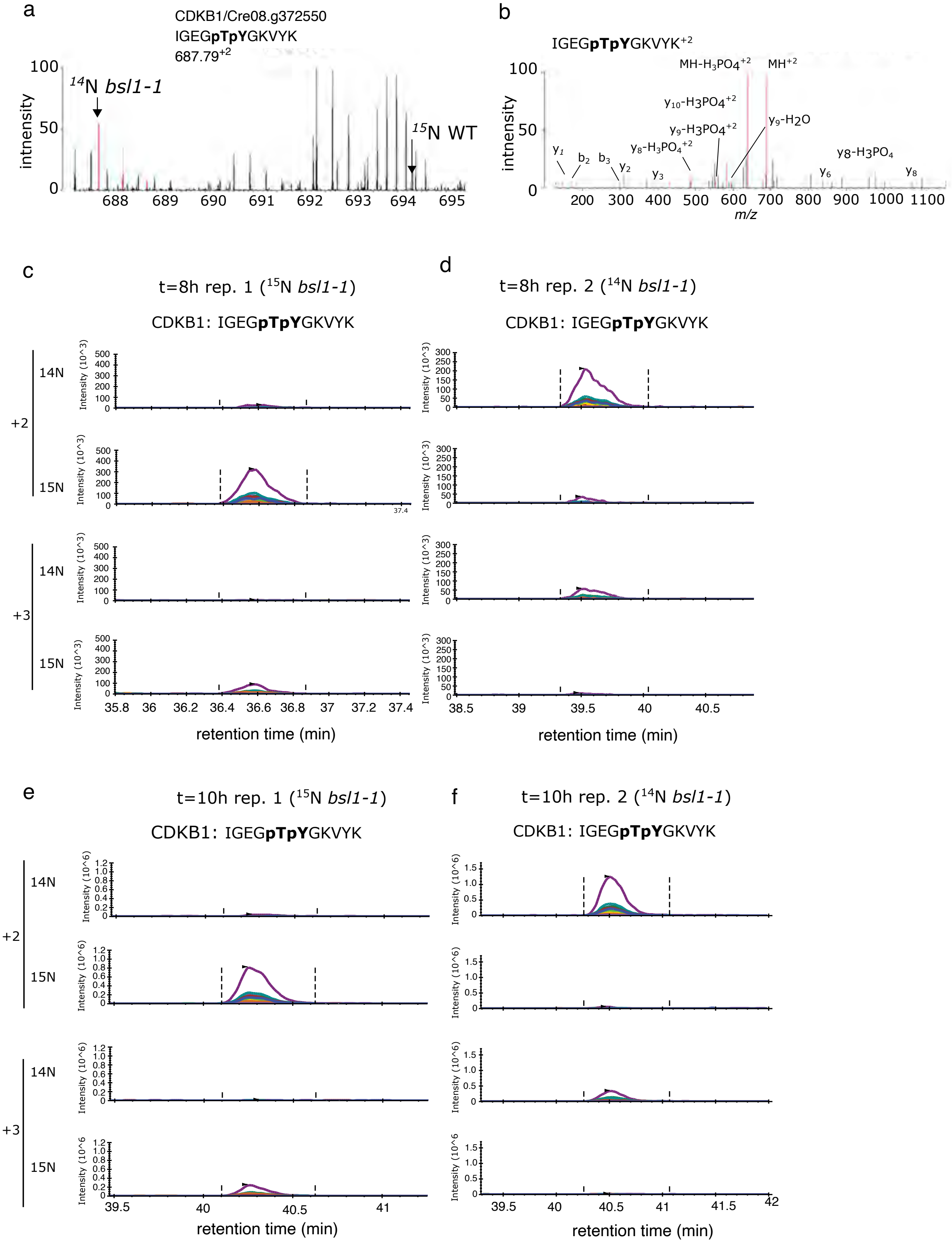


**Extended Data Fig. 6: Targeted quantification of T14,Y15-phosphorylated CDKB1. a**, MS1 spectrum of the doubly phosphorylated CDKB IGEGpTpYGK peptide shows a stronger signal in bsl1-1 (14N) than WT (15N). **b**, MS2 spectrum shows the fragmentation pattern of the m/z=687.79 peak in (a). **(c-f)**, Targeted quantification of the IGEGpTpYGK peptide (phosphorylated on T14 and Y15, with +2 or +3 charge state) in PRM (parallel reaction monitoring) mode. Each colored curve represents the intensity of a specific MS2 ion at a given retention time. Peptides from t=8h and t=10h timepoints were analyzed. The *bsl1-1* mutant cells were metabolically labeled with heavy nitrogen (15N) in rep. 1. In rep. 2, the wild type was labeled with 15N. **c**, t=8h, repeat 1; **d**, t=8h, repeat 2; **e**, t=10h, repeat 1; **f**, t=10h, repeat 2.

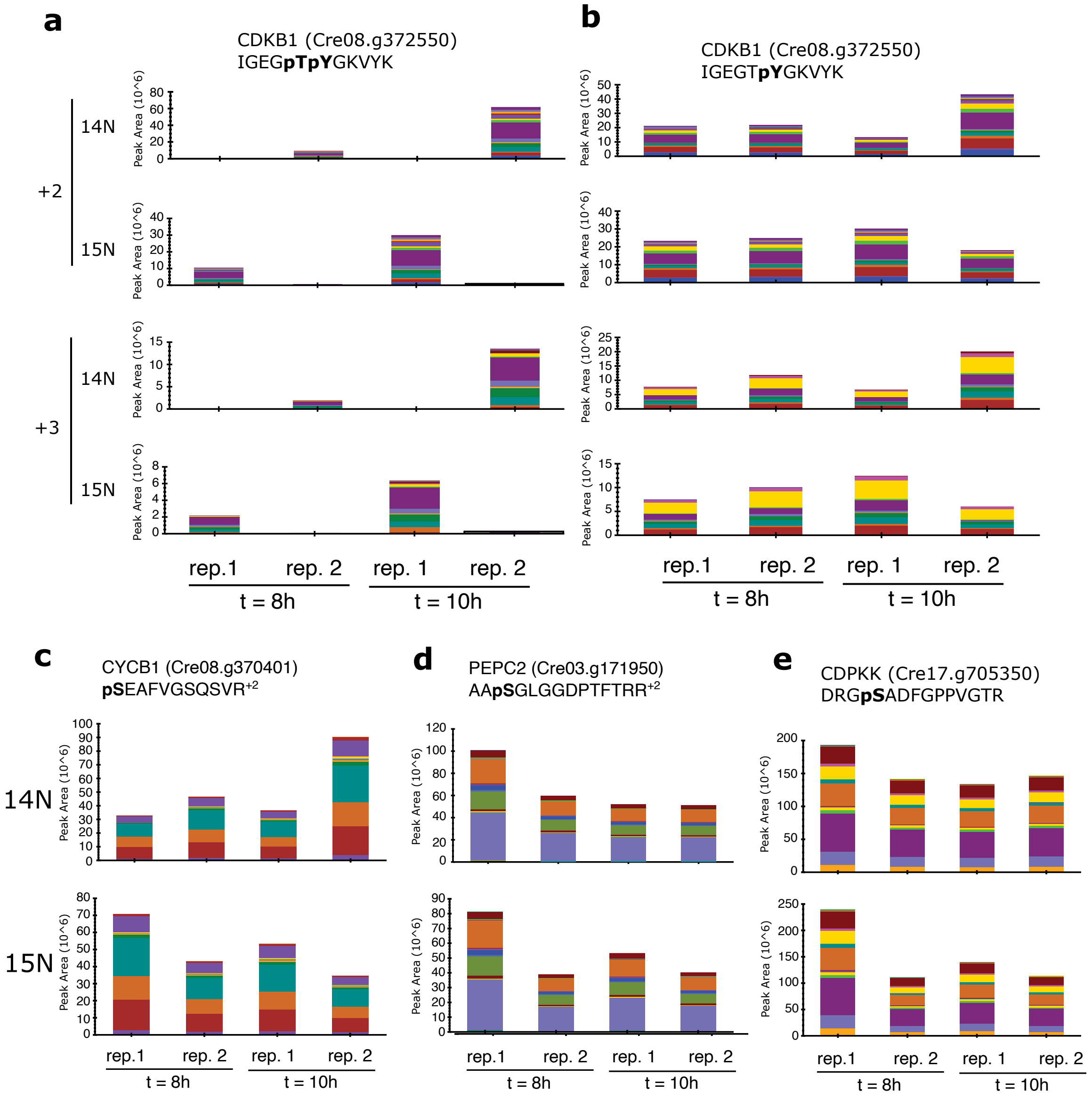


**Extended Data Fig. 7: Summary of PRM MS2 quantification of CDKB1 and control phosphopeptides in *bsl1-1* and wild type.**

**a-e**, Each bar shows the total integrated peak intensity (MS2 quantification, parallel reaction monitoring mode, PRM) of the indicated phosphopeptides. Colors represent the integrated area of different MS2 ions across the elution peak. In all panels, rep.1 = 14N-WT vs 15N-*bsl1-1*, rep.2 = 14N-*bsl1-1* vs 15N-WT. **a**, The doubly phosphorylated peptide IGEGpTpYGK is increased in *bsl1-1* cells compared to wild type (15N/*bsl1-1* peak stronger in rep.1 and 14N/*bsl1-1* peak stronger in rep.2). Ion intensity as a function of retention time for this peptide is shown in fig. S6. **b**, The singly phosphorylated peptide IGEGTpYGK shows similar total integrated intensity in *bsl1-1* and wild type. **c**, Quantification of phosphorylated CYCB1. **d-e**, Quantification of two phosphorylated control peptides, which show similar integrated intensity in *bsl1-1* and wild type at t=8h and t=10h.

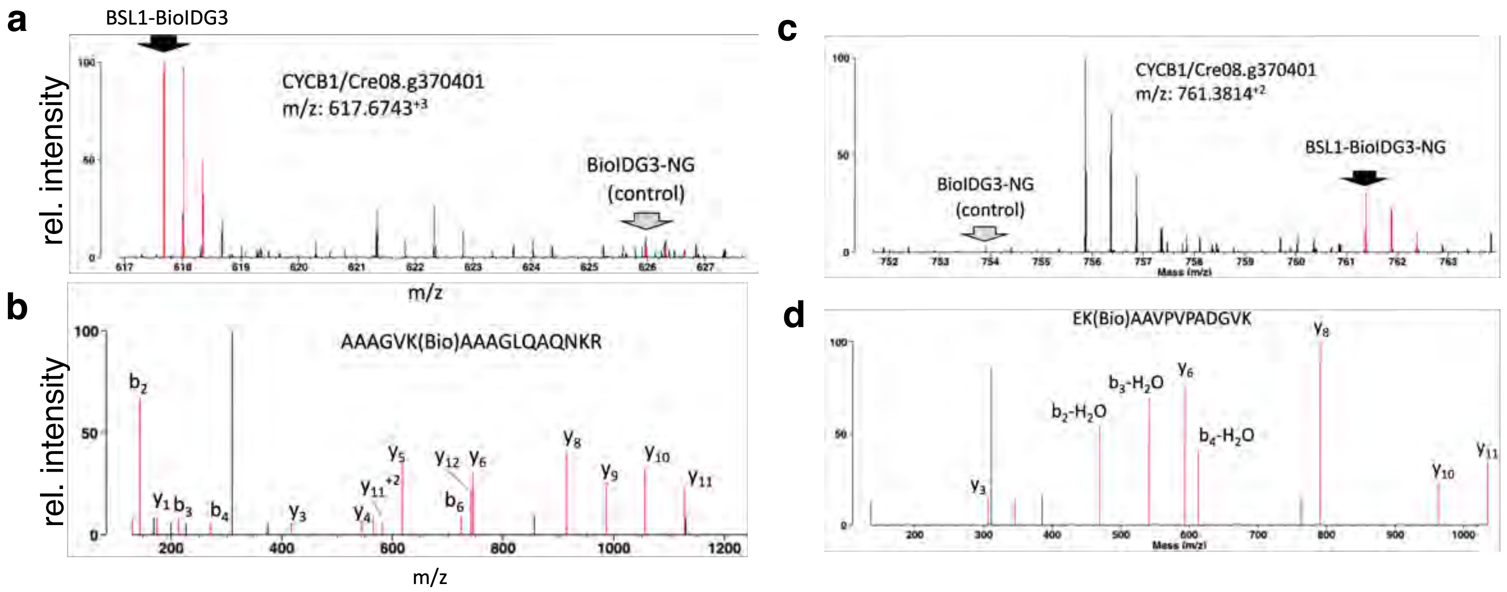


e

intensity ratios of CYCB1 peptides

| peptide | ratio (BSL1/control)[a] | exp.[b] | 15N label[c] |
|---|---|---|---|
| AAAGVK(Bio)AAAGLQAQNKR | 14.4 | 1 | control |
| EK(Bio)AAVPVPADGVK | 8.1 | 1 | control |
| AVQPAEAENGVLGNNNVAK(Bio)AAAGVK | 10.2 | 1 | control |
| AAAGVK(Bio)AAAGLQAQNK | 11.4 | 1 | control |
| EK(Bio)AAVPVPADGVK | 6.9 | 1 | control |
| AVQPAEAENGVLGNNNVAK(Bio)AAAGVK | 1.6 | 2 | BSL1 |
| EK(Bio)AAVPVPADGVK | 9.2 | 3 | BSL1 |
| AAAGVK(Bio)AAAGLQAQNKR | 7.9 | 3 | BSL1 |
| EK(Bio)AAVPVPADGVK | 6.6 | 4 | control |
| AAAGVK(Bio)AAAGLQAQNK | 4.6 | 4 | control |

a: MS1 intensity ratio as reported by Protein Prospector
b: Numbers indicate independent experiments.
c: Indicates whether cells expressing BioIDG3-NG (control) or BSL1-BioIDG3-NG (BSL1) was labeled with heavy $^{15}N$.

**Extended Data Fig. 8. Quantitative proximity labeling indicates proximity between CYCB1 and BSL1. a-b,** BSL1 was fused to an engineered version of the biotin ligase BioID (BioIDG3-NeonGreen, see Methods). BioIDG3-NeonGreen was used as a control. Biotinylated peptides were purified and analyzed by mass spectrometry. **a-b,** Control cells were metabolically labeled with 15N. **a,** MS1 spectrum (black arrow) of the CYCB1 peptide AAAGVK(Bio)AAAGLQAQNKR. The corresponding peak in the control sample (grey arrow) has lower intensity. **b,** MS2 fragmentation pattern for the peptide in (a). **c-d,** The experiment was repeated with the labeling inverted (heavy BSL1-BioIDG3-NG). **c,** MS1 spectrum of the peptide EK(Bio)AAVPVPADGVK (black arrow), **d,** MS2 fragmentation pattern of the peptide in (c). **e,** Table detailing intensity ratios of all CYCB1 peptides detected in four independent experiments. The median intensity ratio (BSL1/control) is 8.

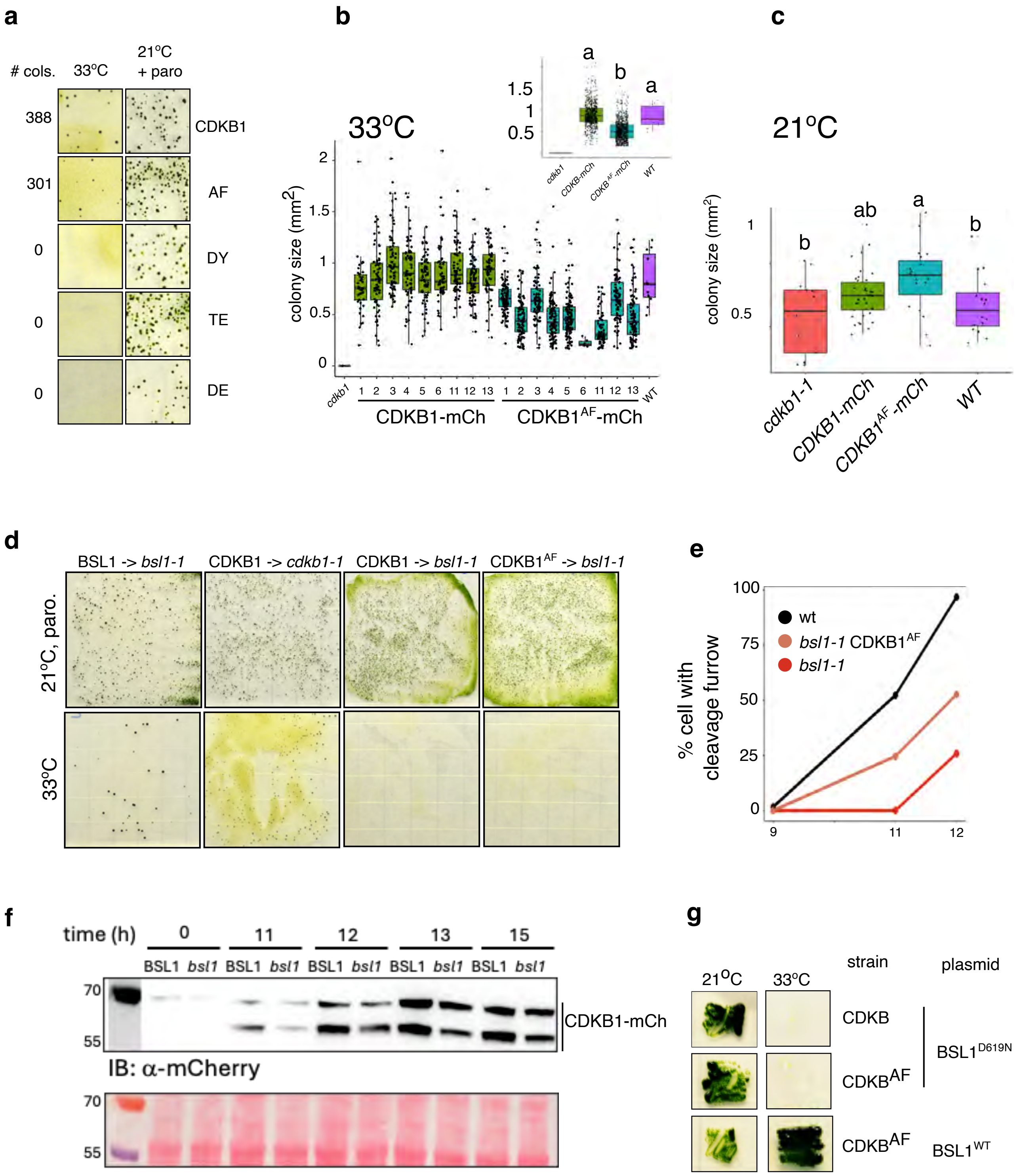


**Extended Data Fig. 9: Phosphorylation of CDKB1 T14 and Y15 is required for optimal growth.**
**a,** The *cdkb1-1* was transformed with wild-type CDKB1-mCherry, and variants containing AF, DY, TE, and DE substitutions at position T14 and Y15. Only wild type CDKB1 and the CDKB1$^{AF}$ variant could rescue *cdkb1-1* at 33oC. Transformation efficiency was assessed by plating half of the transformed cells on plates containing paromomycin at 21oC. **b-c,** Quantification of colony size of individual clones of *cdkb1-1* rescued by CDKB1-mCh and CDKB1$^{AF}$-mCh after 5 days at 33°C **(b)**, and 7 days at 21°C **(c)**. Insert: combined data for CDKB1-mCh and CDKB1AF-mCh. Box plots represent median, 25th/75th percentile with whiskers max 1.5 * inter-quartile range. Letters indicate significance levels, Tukey post-hoc p<0.05. **d**, Complementation experiments. The *bsl1-1* and *cdkb1-1* mutants were transformed with plasmids containing the BSL1 and CDKB1 genes. The *bsl1-1* mutant was rescued by BSL1 but not  by with wild-type CDKB1 or CDKB1$^{AF}$. Transformed cells were split evenly. Half of the cells were plated on 10 µg/ml paromomycin at a permissive temperature (21oC) as a transformation efficiency control. The other half was plated at 33°C to select for complementation of the *bsl1-1* or *cdkb1-1* mutants. **e,** Timing of cleavage furrow formation in *bsl1-1* and *bsl1-1* CDKB$^{AF}$. Each data point represents data from two independent experiments. 20-30 cells were counted in each experiment. **f**, α-mCh western blot of CDKB1$^{AF}$-mCh in synchronized cells (BSL1 +/. BSL1-mCh runs as two bands (*, and **). **g**, *bsl1-1* CDKB1 and *bsl1-1* CDKB$^{AF}$ cells were transformed with phosphatase-dead BSL1$^{D619N}$-Venus or BSL1$^{WT}$-Venus, and selected on paromomycin at 21°C. Expression of BSL1 was verified by localization to spindle poles. Rescue of *bsl1-1* was tested at 33°C. Only BSL1$^{WT}$ rescued *bsl1-1*.

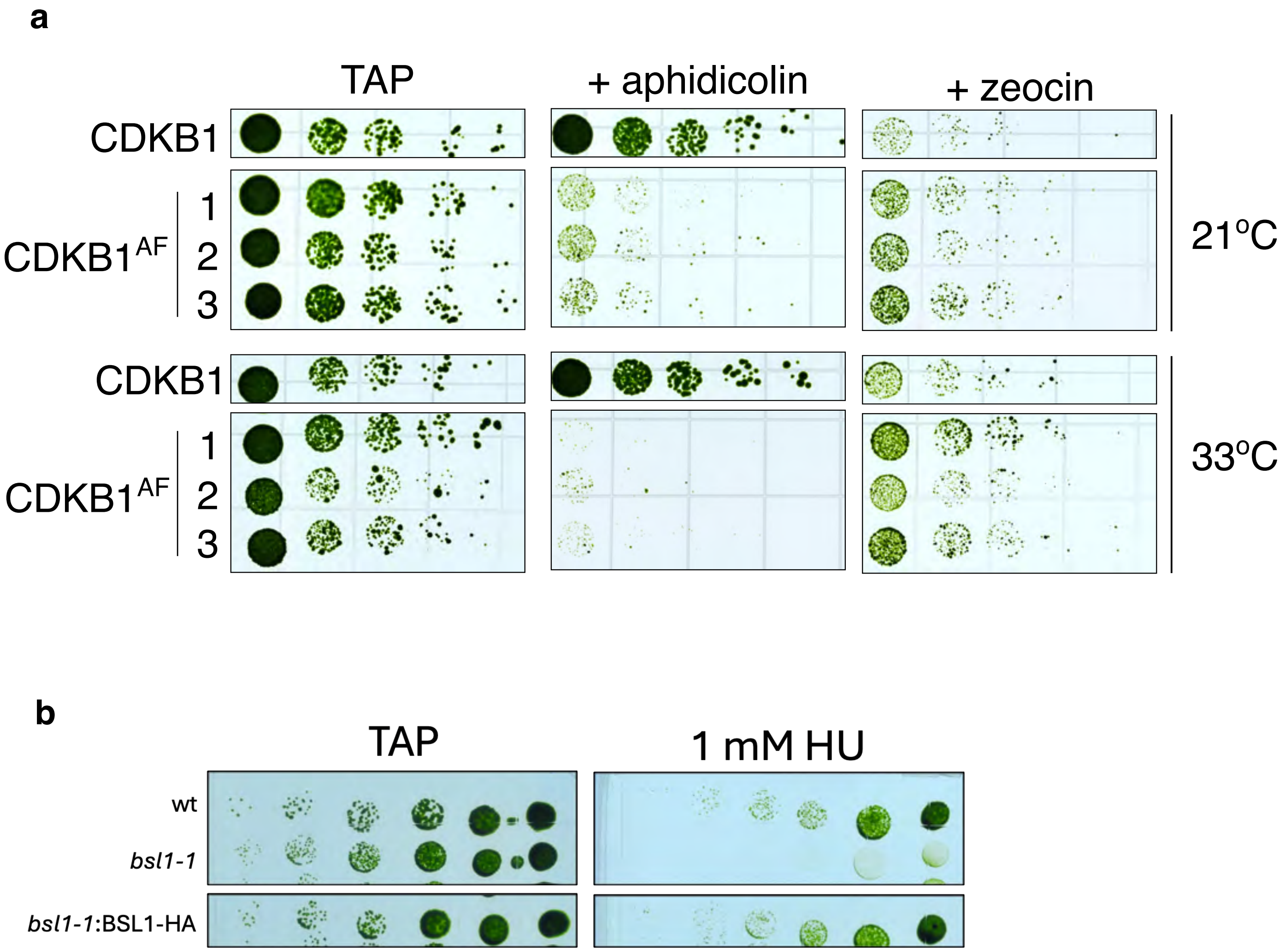


**Extended Data Fig. 10. Mutations of CDKB1-T14/Y15 and BSL1 alter the sensitivity to DNA replication stress. a,** Cells expressing $CDKB1^{AF}$ are hypersensitive to the polymerase inhibitor aphidicholin, but not to the DNA-damage-inducing drug zeocin. *cdkb1-1* cells complemented with either wild type CDKB1-mCherry (CDKB1) or unphosphorylatable $CDKB1^{AF}$-mCherry ($CDKB1^{AF}$, three independent clones). The density of each culture was normalized to $OD_{750}$ ~0.4 and cells were spotted in 5-fold serial dilutions on TAP agar plates containing 50 µM aphidicholin or 2 µg/ml zeocin and incubated at 21°C and 33°C. **b,** Sensitivity of the *bsl1-1* mutant to HU. Cells were spotted (5-fold serial dilutions) on TAP agar plates, and plates supplemented with 1 mM HU, and incubated at at 21°C.

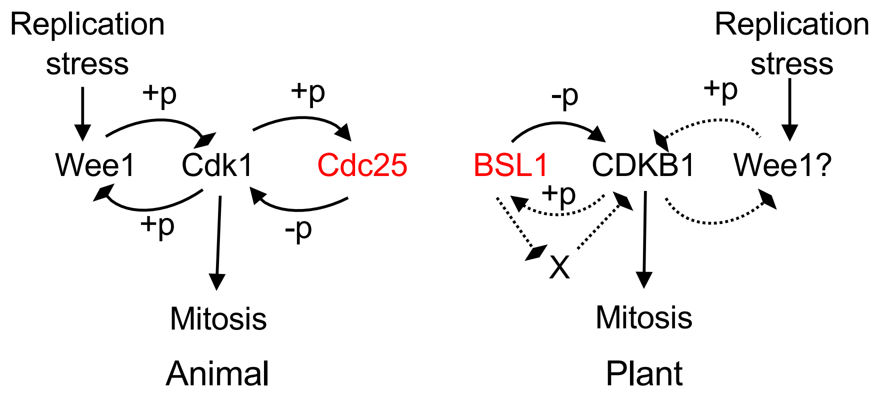


**Extended Data Fig. 11. Proposed model for the role of BSL1 and CDKB1 in control of mitosis in *Chlamydomonas* compared to animals.** In fungi and metazoans, mitosis is triggered by the activation of Cdk1, which is inactivated during replication stress by phosphorylation at T14/Y15 and subsequently activated by Cdc25-mediated dephosphorylation. Similarly, in Chlamydomonas, CDKB1 is essential for mitosis and is inactivated by phosphorylation at T14/Y15. However, dephosphorylation of CDKB1 T14/Y15 is mediated by BSL1. In addition, BSL1 promotes CDKB1 accumulation independent of T14/Y15 dephosphorylation, possibly by inhibiting an unknown protein (X) that mediates CDKB degradation. +p/-p: phosphorylation/dephosphorylation. Triangle arrowheads: positive regulation. Diamond arrowheads: inhibitory regulation. Dashed lines: hypothetical mechanisms.
